# Modeling steady state thermoregulation of near-term human fetus

**DOI:** 10.64898/2026.08.13.744721

**Authors:** Alyssa Payne, Ankit Joshi, Shri H. Viswanathan, Savan P. Shah, Daibo Zhang, Stephanie E. Lindsey, Konrad Rykaczewski

## Abstract

Maternal thermal strain is associated with adverse pregnancy outcomes, yet fetal temperatures cannot currently be directly measured, limiting quantification of fetal thermal strain. Here, we develop two steady-state models for estimating internal temperatures in a near-term fetus. First, we improve the only previously published human fetal thermoregulation model, deriving a closed-form solution within its simplified uniform-cylinder representation. Second, we introduce a multilayer, anatomically segmented model that resolves tissue-specific temperatures. Both couple the fetal body to central blood pool and amniotic fluid compartments and incorporate a new placenta–umbilical cord heat-exchanger representation. Predictions agree with available intrauterine scalp measurements, with fetal core and head-center temperatures approximately 0.5°C and 0.8°C above maternal core, respectively. Physiologically plausible changes in umbilical cord heat-exchanger effectiveness or blood flow increased fetal temperatures by approximately 0.3°C. These models enable estimation of otherwise inaccessible temperatures, while the multilayer formulation lays a foundation for transient, coupled maternal-fetal thermoregulation modeling.

## 1. Introduction

Maternal thermal strain resulting from heat or cold exposure, high exertion, or fever has been associated with preterm birth, low birth weight, congenital anomalies, maternal hypertension, and gestational diabetes [1–3]. However, the quantitative relationship between measurable thermophysiological parameters and fetal temperature remains poorly understood [1,3]. Human intrauterine measurements have been limited to scalp and amniotic fluid temperatures near delivery [4–8]. Consequently, fetal aortic and esophageal measurements from lambs and baboons [9–12] were used to formulate the now standardized assumption that fetal core temperature is approximately 0.5°C warmer than the maternal core. This assumption, together with maternal physiological responses measured during controlled heat-stress trials, has informed safety guidelines for heat exposure and exercise during pregnancy [13,14]. However, such recommendations warrant caution in the absence of direct measurements of human fetal core temperature, which may eventually become feasible through emerging technologies [15]. Moreover, it remains unclear how temperatures across different fetal anatomical regions change with increasing maternal core temperature. A computational model of fetal thermoregulation can quantify the relationship between maternal and fetal temperatures, including at experimentally inaccessible anatomical locations.

Compared with adults, the fetus has two-fold higher mass-specific metabolic heat generation rate, lacks sweating and shivering, and dissipates heat primarily through umbilical-placental blood flow (∼85%), with the remainder transferred by convection to the amniotic fluid (∼15%) [3]. The only human fetal thermoregulation model, described by Guenkawa et al. [16] and based on an earlier ovine simulation [17], represents the fetus as uniform head, torso, leg, and arm cylinders, neglecting tissue distributions. The model is also inherently restricted to steady-state formulation and lacks basic parameter values needed for replication.

To address these limitations, we developed two fetal thermoregulation models: an improved closed-form solution for the simplified uniform cylinder representation and a more anatomically realistic model adapted from our improved adult Stolwijk model [18,19]. The latter enables estimation of tissue-specific temperatures by representing the head, torso, arms, legs, hands, and feet, with each segment divided into concentric core (viscera and bone), muscle, fat, and skin layers. Both models represent a near-term (39-week, 3.2 kg [20]) fetal body coupled to well-mixed arterial blood and amniotic fluid compartments and incorporate new two-heat-exchanger submodel to account for umbilicoplacental heat transport [21–23] (see **Figure 1**). We assume steady-state in both models, a reasonable first approximation given the slow rate of change in maternal core temperature. However, the multilayer cylinder model can be readily extended to transient simulations and coupled with a future maternal thermoregulation model. Model parameters, including tissue thermophysical properties and regional perfusion are based on established physiological data and fetal hemodynamic measurements. We describe the formulation and validation of both models and use them to quantify how physiological parameters influence fetal temperature elevation relative to the maternal core.

**Figure 1.**
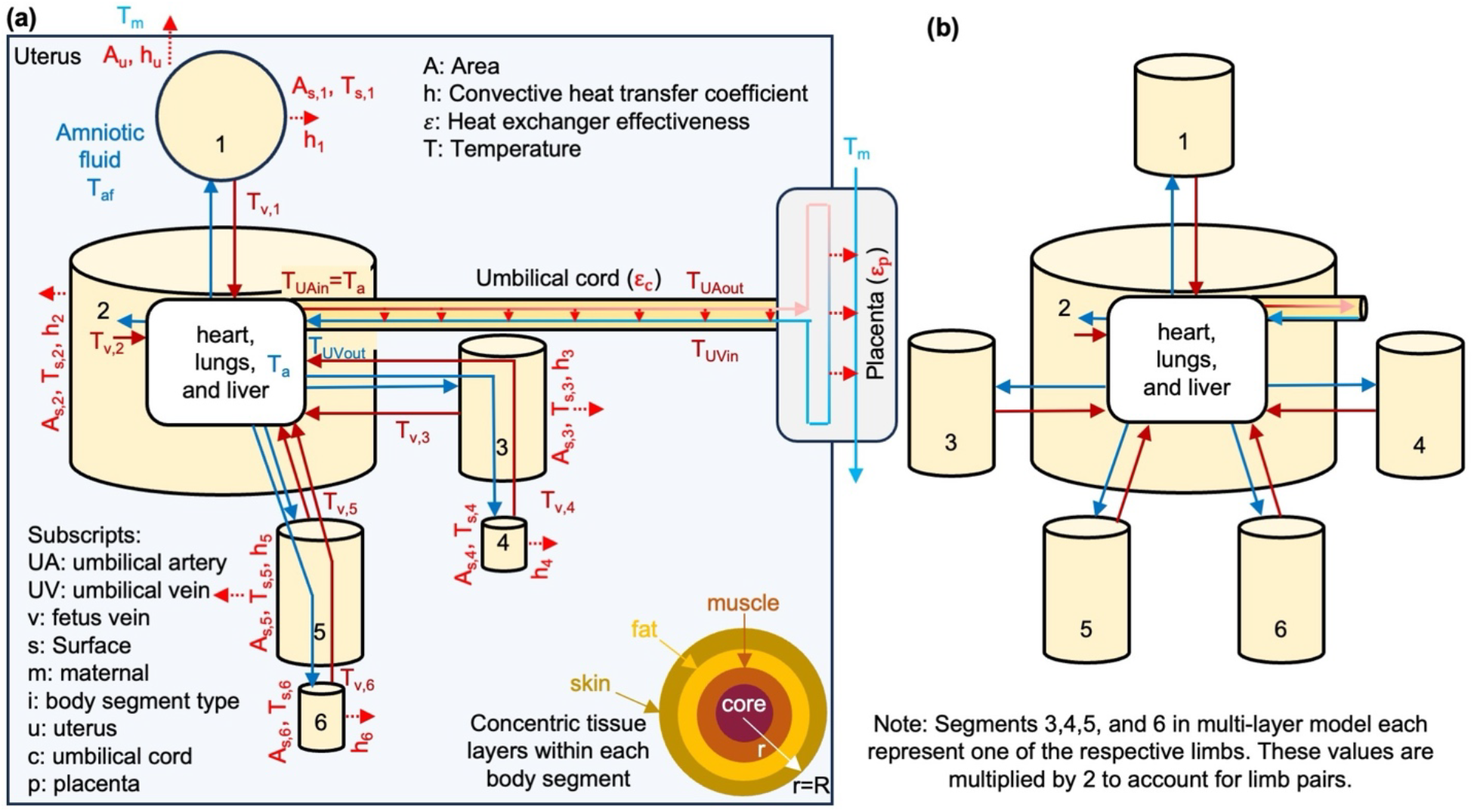
Schematic of the **(a)** uterus with multilayer cylinder fetus, umbilical cord, and placenta models with major blood flow (arrows with solid lines) and heat transfer (arrows with dashed lines) indicated; inset shows a multilayer cross section for each tissue layer (segments 3-6 contain a bilateral multiplier to account for limb pairs) and **(b)** six uniform cylinder fetus.

### 2. Methods: overview of the models

Both models include four subsystems: (i) placenta-umbilical cord coupled heat exchangers, (ii) amniotic fluid, (iii) central arterial blood pool, and (iv) fetal body. The umbilical cord consists of two arteries that are coiled around a single central vein, all embedded within a cylinder made of Wharton’s Jelly [22], and is modeled as an insulated countercurrent heat exchanger characterized by its effectiveness, ε_c_ (i.e., the ratio of actual to maximum heat transfer in-between the fluids occurring in the heat exchanger). The placenta is modeled as a shell-and-tube heat exchanger [19,23], characterized by very high effectiveness, ε_p_, based on animal experiments [13]. The umbilical-venous temperature relationship is derived in the Supplementary Material (SM).

We formulate heat-rate balances for the central blood pool and the amniotic fluid. In the uniform cylinder model, we use solutions to steady-state radial Penne’s bioheat transfer equation and heat rate balance for each homogeneous cylinder to derive “constitutive equations” for segmental skin and venous temperatures. By combining these equations with the central blood pool and amniotic fluid heat-rate balances, we obtain closed-form solutions for the temperatures of the central blood pool and amniotic fluid as functions of maternal core temperature, body geometry, tissue properties, blood circulation, and the effectiveness of the placenta and umbilical cord. The multilayer model solves the set of linear equations obtained from performing a steady-state heat-rate balance on each of the 24 discrete, uniform tissue compartments, each represented by a single temperature (four tissue layers in each of six body segments), as well as on the central blood pool and amniotic fluid. The model formulation, implementation, and Matlab codes are described in depth in the SM.

## 3. Results and Discussion

Fetal scalp temperatures predicted by both models (**Figure 2a**) agree closely and fall near the center of the range of intrauterine measurements reported by Lavesson et al.[8] during vaginal labor with maternal axillary temperatures of 35.5-39.5°C (assumed to be 0.27°C below maternal core [8]). For fixed physiological inputs, fetal-to-maternal core temperature offsets remain constant for the amniotic fluid (0.58 and 0.43°C for uniform and multi-layer models, respectively), scalp (0.53 and 0.46°C), core (i.e., central blood pool; 0.48 and 0.52°C), and center of the head (0.76 and 0.78°C). Although experimental studies vary in measurement location, protocol, and sample size [4–8], model predictions are consistent with the observed small (≤0.2°C) skin-to-amniotic-fluid or uterine-wall temperature differences and larger (0.3–0.5°C) scalp-to-maternal-core differences.

Following validation, we used the models to analyze internal fetal temperatures that are not accessible through human measurements. Models show some appreciable differences in internal temperature distributions among body segments, while all skin temperatures converge towards the amniotic fluid temperature (**Figure 2b**). The head and torso exhibit the largest center-to-skin temperature differences, 0.32°C and 0.14°C, respectively. These segments have the highest heat generation within the fetal body, partially offset by high blood perfusion that moderates the resulting center-to-skin temperature gradients. In contrast, the center-to-skin temperature differences in the arms, legs, hands, and feet are only ∼0.075°C. Consistent with fetal physiology, the fetal “core,” represented by the central blood pool, remains close to the torso temperature, differing by ≤0.1°C.

**Figure 1.**
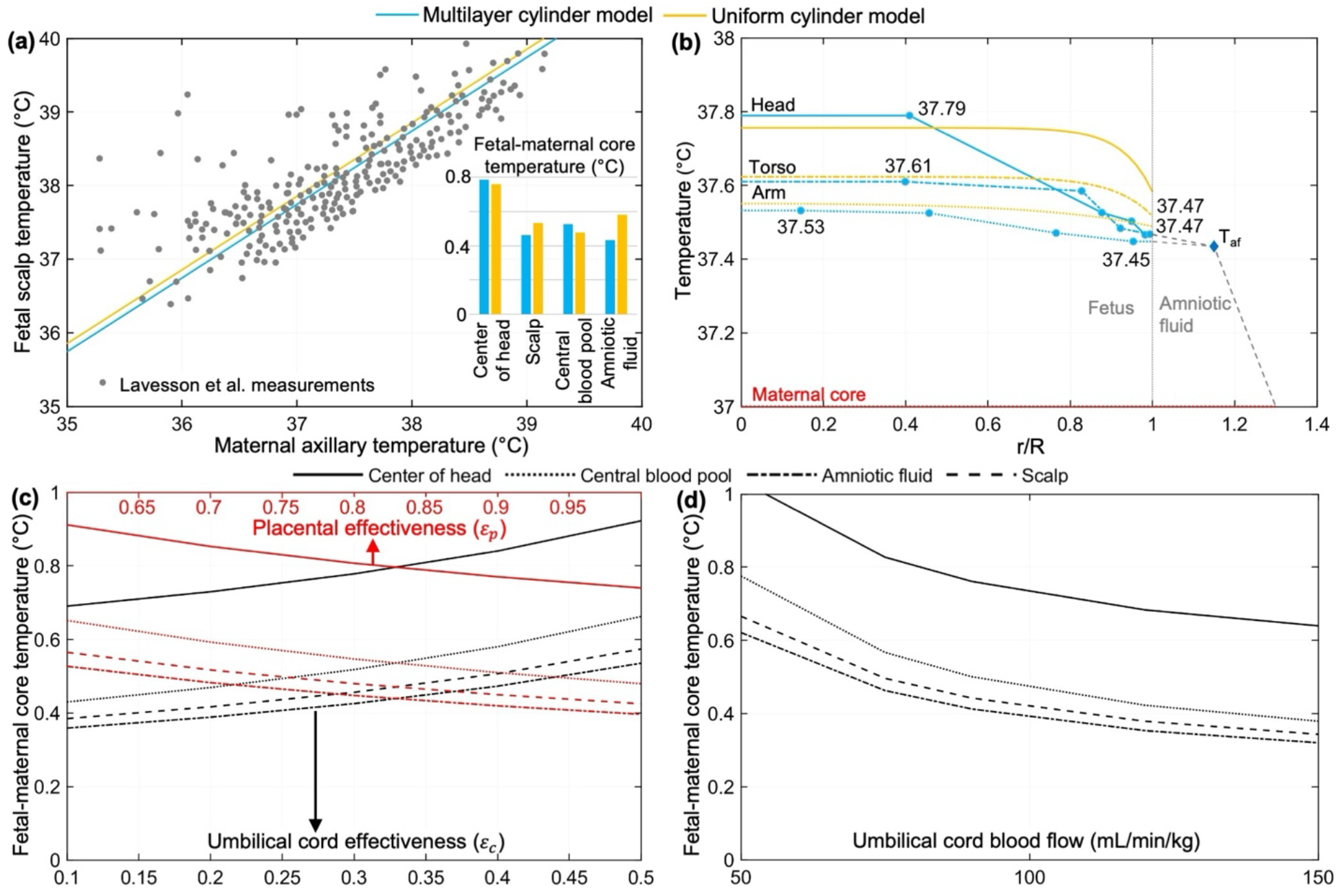
**(a)** Comparison of predicted fetal scalp and maternal axillary temperatures against Lavesson et al. [8] (inset: offsets of scalp, amniotic fluid, central blood pool, and center of head temperatures from maternal core); **(b)** predicted radial temperature variation in head, torso, and arm as function of scaled radius (r/R) (hand, leg, and feet variation are comparable to arm and shown in SM); **(c-d)** offset of fetal temperatures from maternal core predicted by multi-layer model with variation in **(c)** placenta and umbilical cord effectiveness **(d)** the umbilical vein flow normalized by fetal weight for multilayer model.

At the fetal surface, the small temperature differences between the skin and amniotic fluid can be explained by the high convective heat transfer coefficients associated with liquid immersion, which range from approximately 200 to 550 W/m^2^°C (see SM, similar values are obtained using free-convection correlations [16]). A simple scaling analysis illustrates the resulting strong thermal coupling. For a fetal surface area of ∼0.2 m^2^, with approximately 20% of the total 8 W metabolic heat generation dissipated through the skin, the average skin heat flux is ∼8 W/m^2^. Therefore, the expected skin-to-amniotic-fluid temperature differences are ΔT∼q″/h∼8/ (200–550) ∼0.015-0.04°C. The uniform model predicts a larger scalp-to-amniotic-fluid temperature difference because head has a large fraction of the total metabolic heat generation (4.6 of 8 W) and the convective heat transfer coefficient for its cylindrical head geometry (224 W/m^2^°C) is lower than that for the spherical head in the multilayer model (306 W/m^2^°C).

Deviations of ε_p_ and ε_c_ from their baseline values of 0.85 and 0.32, respectively, can substantially affect fetal temperatures (**Figure 2c**). As ε_p_ →1 the placenta approaches a perfect heat sink, cooling fetal blood toward maternal temperature and thereby producing the lowest achievable fetal temperatures, approximately 0.1°C below their baseline values relative to maternal core temperature. In contrast, reducing ε_p_ from 0.85 to 0.6 to simulate the potential restriction of maternal blood supply to the placenta during heat strain [3] increases all fetal temperatures by 0.25°C relative to baseline. The effect of ε_c_ is reversed because the umbilical cord acts as a conduit carrying blood to and from the placental heat sink. Increasing ε_c_ from 0.32 to 0.5, as could potentially occur if hypercoiling increases the heat-exchange area between the two blood streams [22], increases fetal temperatures by 0.15°C relative to baseline. In the extreme limiting case of ε_c_ →1, most of the excess heat carried by the umbilical arteries is transferred directly to the returning umbilical venous blood before reaching the placenta, resulting in fetal temperature increases of more than 2°C and potentially raising brain temperature to ∼40°C, a range associated with teratogenic risk [2]. Reducing umbilical blood flow from the baseline value of 100 mL/min/kg to 50 mL/min/kg, the lower end of the reported physiological range associated with hypercoiling [24], increases fetal temperatures by 0.25°C (**Figure 2d**). Under these conditions, the fetal “core” and “brain” exceed maternal core temperature by about 0.8 and 1.1°C, respectively, substantially exceeding the generally assumed 0.5°C offset.

## 5. Conclusions

In this work, we provide two approaches to quantifying the relationship between maternal and fetal temperatures by improving the only previously published human fetal thermoregulation model, which uses a simplified uniform-cylinder formulation, and introducing a more anatomically detailed model that resolves multiple concentric tissue layers within cylindrical and spherical representations of fetal body segments. Despite their different levels of anatomical detail, the uniform-cylinder and multilayer models produced similar predictions and agreed with available fetal scalp and intrauterine temperature measurements. The models predict that in baseline physiological conditions fetal skin remains near the amniotic fluid temperature, whereas the deep torso and head remain modestly warmer (0.15 to 0.3°C). The models further predict that physiologically plausible reductions in umbilical blood flow from 100 to 50 mL/min/kg can increase fetal core and head-center temperatures by approximately 0.3°C, quantifying the thermal impact of changes in umbilical cord circulation.

While the uniform-cylinder model provides a simple approach for estimating maximum fetal temperatures under quasi-steady-state conditions, the multilayer model establishes a flexible foundation for future fetal thermoregulation modeling. The latter can be readily extended to transient conditions, coupled with a maternal thermoregulation model developed by adapting an adult female model using an equivalent compartmental formulation, and adjusted to reflect anatomical and physiological changes throughout gestation. The models can be further refined using multiphysics simulations that more directly link placental and umbilical heat-exchange effectiveness to underlying physiological parameters. Such an integrated framework could support the evaluation of fetal thermal strain during environmental heat exposure, exercise, pregnancy complications, and fetal surgical interventions, while informing thermal management strategies and future guidelines for heat exposure during pregnancy.

