## Supplemental Materials for "Modeling steady state thermoregulation of near-term human fetus"

**Supplemental Material for "Modeling steady state thermoregulation of near-term human fetus"**

Alyssa Payne,[1†] Ankit Joshi,[1,2†] Shri H. Viswanathan,[1] Lindsey, S.,[3,4] Daibo Zhang,[3] Savan P. Shah,[3] and Konrad Rykaczewski [1,2*]


| Section | Section Name | Page |
|---|---|---|
| S.1 | The umbilical cord and placental heat exchange | 2 |
| S.1.2 | The placenta | 2 |
| S.1.3 | The umbilical cord | 2 |
| S.1.4 | Combined umbilical cord and placental heat exchange | 3 |
| S.2 | Fetal skin-to-amniotic fluid convection and uterine wall heat transfer | 3 |
| S.2.1 | Convective heat transfer of the fetal skin | 3 |
| S.2.2 | Heat transfer between the amniotic fluid, uterine wall, and maternal core | 4 |
| S.2.2.1 | Convection between the uterine wall and the amniotic fluid | 4 |
| S.2.2.2 | Conduction through amnion and chorion layers | 5 |
| S.2.2.3 | The maternal side "heat sink" limited by the myometrial blood perfusion | 5 |
| S.3 | Formulation and implementation of the two fetal thermoregulation models | 6 |
| S.3.1 | Uniform cylinder model formulation and implementation | 6 |
| S.3.2 | Multi-compartmental cylinder model formulation and implementation | 9 |
| S.4 | From fetal anatomical to cylinder (and sphere) fetal geometry | 9 |
| S.4.1 | The head geometry | 10 |
| S.4.2 | The trunk geometry | 10 |
| S.4.3 | The limb geometries | 10 |
| S.5 | The blood flow rate and distribution within the fetus | 12 |
| S.6 | Thermal conductivity and inter-layer conductance of the fetal tissues | 13 |
| S.7 | Metabolic heat generation of the fetal tissues | 15 |
| S.8 | Radial temperature variation in hands, legs, and feet | 17 |
| S.9 | MATLAB code implementation of uniform model | 18 |
| S.10 | MATLAB code implementation of multilayer cylinder model | 20 |
| S.11 | Lavesson et al. maternal axillary vs. fetal scalp temperatures for model validation | 38 |
| — | References | 45 |

### S.1 The umbilical cord and placental heat exchange

Accurate modeling of placental and umbilical transport is particularly important because approximately 85% of fetal heat transfer to the mother is estimated to occur through placental circulation rather than through fetal skin and the uterine wall [1–3]. However, the coupled blood flow and transport processes within the placenta and umbilical cord are highly complex [4–6], making direct incorporation of detailed simulations into whole-body thermoregulation models computationally impractical. The placenta and umbilical cord are examples of biological systems that can be modeled as heat exchangers. The umbilical cord consists of one vein and two arteries that are coiled around the vein. This geometry aligns with that of a countercurrent heat exchanger, and the umbilical cord can be modeled as such. In many flow conditions, the heat transfer between the maternal and fetal sides of the placenta can be modeled as a concurrent-flow tube-and-shell heat exchanger [7,8]. Next, we describe the heat exchanger models for the placenta, umbilical cord, and their coupled system, which relate maternal core temperature to the inlet of the umbilical cord arteries and the outlet of the umbilical cord vein.

#### S.1.2 The placenta

The placenta sits between two regions, the amniotic fluid and the uterine wall. Within the placenta, heat transfer occurs between fetal and maternal blood, but the two streams remain separated. Maternal blood is pumped into the placenta, passing over the outside of the villous trees within the intervillous space. The terminal villi are capillaries with a thin-walled membrane through which nutrients, oxygen, carbon dioxide, and heat diffuse between maternal and the warmer fetal blood that perfuses these microchannels. After fetal blood returns through the placenta and umbilical vein, the fetus absorbs oxygen and nutrients. Considering the flow of maternal blood over the fetal blood-filled vessel network, the placenta is often modeled as a shell-and-tube heat exchanger. The heat-exchanger effectiveness of the placenta is estimated at 0.85 [9], based on prior animal studies. Also, taking into consideration that near-term uteroplacental (maternal side) blood flow substantially exceeds fetoplacental blood flow, the placental heat exchanger effectiveness ($\varepsilon_p$) may be expressed as a function of maternal core temperature ($T_m$), outlet temperature of the umbilical arteries ($T_{UA,out}$), and inlet temperature to the umbilical cord vein ($T_{UV,in}$) [10]:

$$\varepsilon_p = (T_{UA,out} - T_{UV,in})/(T_{UA,out} - T_m) \tag{S1}$$

which can be rearranged to:

$$T_{UV,in} = T_{UA,out}(1 - \varepsilon_p) + \varepsilon_p T_m \tag{S2}$$

We need to obtain a relationship for the temperature of the blood returning to the fetus from the placenta ($T_{UV,out}$) as a function of the temperature of the blood leaving the fetus ($T_{UA,in}$), the $T_m$, and the characteristics of the two heat exchangers ($\varepsilon_c$ and $\varepsilon_p$).

#### S.1.3 The umbilical cord

Assuming that the cord is insulated from the amniotic fluid by Wharton's jelly and that the blood flow rate in the umbilical vein and arteries are equal, heat balance on the insulated cord reduces to [10]:

$$T_{UA,in} - T_{UA,out} = T_{UVout} - T_{UV,in} \tag{S3}$$

From the definition of the effectiveness, we obtain:

$$\varepsilon_c = \frac{m_{AV} c_b (T_{UA,in} - T_{UA,out})}{m_{UV} c_b (T_{UA,in} - T_{UVin})} \rightarrow T_{UA,out} = T_{UA,in}(1 - \varepsilon_c) + \varepsilon_c T_{UVin} \tag{S4}$$

Then finally substituting Eq. S4 into Eq. S8, we obtain:

$$T_{UV,out} = T_{UV,in}(1 - \varepsilon_c) + T_{UA,in}\varepsilon_c \quad (S5)$$

The physical constants utilized in both the multi-layer and uniform cylinder models are the mass flow rate of blood through the umbilical artery, blood density ($\rho_b$ = 1050 kg/m$^3$), blood flow through the umbilical vein ($Q_{UV}$ = 318 mL/min [11]), the weight of the fetus (3.2 kg), and specific heat capacity of blood (3617 J/kgK). The mass flow rate of blood through the umbilical cord artery is determined by:

$$m_{UA} = \rho_b * Q_{UV} * 10^{-6} * \frac{\mathrm{Weight}_{fetus}}{60\ \mathrm{s/min}} = 5.62 * 10^{-3}\ \mathrm{kg/s} \quad (S6)$$

#### S.1.4 Combined umbilical cord and placental heat exchange

We aim to obtain a relationship for the temperature of the blood returning to the fetus from the placenta ($T_{UV,out}$) as a function of the temperature of the blood leaving the fetus ($T_{UA,in}$), the $T_m$, and the characteristics of the two heat exchangers ($\varepsilon_c$ and $\varepsilon_p$). Substituting rearranged Eq. S5 into S2, we obtain:

$$T_{UV,in} = \frac{T_{UA,in}(1 - \varepsilon_p)(1 - \varepsilon_c) + T_m\varepsilon_p}{1 - \varepsilon_c(1 - \varepsilon_p)} \quad (S7)$$

Then, finally substituting the above equation into Eq. S5, we obtain:

$$T_{UV,out} = T_{UA,in}\left(\varepsilon_c + \frac{(1-\varepsilon_p)(1-\varepsilon_c)^2}{1-\varepsilon_c(1-\varepsilon_p)}\right) + T_m\left(\frac{\varepsilon_p(1-\varepsilon_c)}{1-\varepsilon_c(1-\varepsilon_p)}\right) = T_a Z_5 + T_m Z_6 \quad (S8)$$

### S.2 Fetal skin-to-amniotic fluid convection and uterine wall heat transfer

In steady state, the amniotic fluid compartment is governed by an energy balance in which heat received from all fetal skin surfaces equals heat dissipated through the uterine wall, as given by Eq.S9. Convective heat transfer from each fetal skin surface to the amniotic fluid is characterized by segment-specific coefficients, which are calculated using a correlation for a cylinder (for every body part except the head) or sphere in crossflow (for the head) of water at 37°C (amniotic fluid approximation) moving at a representative speed of the fetal limbs of 1 cm/s (see rationale for this value below). The convective heat exchange $h_{sk,i} \cdot A_{s,i}$ couples each skin node to the amniotic fluid compartment.

$$0 = \sum_{i=1}^{6} \underbrace{h_{sk,i}\, A_{s,i}\, (T_{sk,i} - T_{af})}_{\text{heat from fetal skin to amniotic fluid}} - \underbrace{h_u\, A_u\, (T_{af} - T_m)}_{\text{heat from amniotic fluid to uterine wall}} \quad (S9)$$

Where, $h_{sk,i}$ is convective heat transfer coefficient at the skin surface of segment i [W m$^{-2}$ K$^{-1}$], $A_{s,i}$ is skin surface area of segment i [m$^2$], $T_{sk,i}$ is skin-layer temperature of segment i [°C], $T_{af}$ is amniotic fluid temperature [°C], $h_u$ is 22 W m$^{-2}$ K$^{-1}$ effective uterine wall heat transfer coefficient, $A_u$ is 0.15 m$^2$ non-placental inner uterine wall area (see further details below).

#### S.2.1 Convective heat transfer of the fetal skin

The convective heat transfer coefficients, $h_i$, of the fetal skin were calculated utilizing the outer diameter of each body segment and utilizing the Hilpert correlation for crossflow of a cylinder

[12] and the Whitaker correlation for crossflow of a sphere [13] (only used for the multi-layer segment model head). The amniotic fluid is approximated as water at 37 °C with a velocity of 1 cm/s (rationale for this representative speed is detailed below). With these assumptions, the relevant values for the amniotic fluid are: Prandtl, Pr, is 6.13, density, ρ, is 1,000 kg/m$^3$, viscosity, μ, is $6.9 * 10^{-4}$ Pa $*$ s and thermal conductivity, $k_{af}$, is 0.6 W/mK. The Hilpert correlation for a cylinder in crossflow:

$$Nu_D = CRe_D^m Pr^{\frac{1}{3}} \quad (S10)$$

Since the Reynolds number in each scenario is between 4 and 4,000, C=0.683 and m=0.466.
Whitaker correlation for a sphere in crossflow:

$$Nu_D = 2 + \left[0.4Re^{1/2} + 0.06Re^{2/3}\right]Pr^{0.4} \quad (S11)$$

The convective heat transfer coefficient can now be derived for each segment (see Table 1 below for values):

$$h_i = \frac{Nu_D k}{D} \quad (S12)$$

Both models have the same volume for each body segment. In the uniform model, the head is a cylinder; in the multi-layer model, it is a sphere. To calculate the head dimensions for the uniform model, its volume was set to that of the head in the multi-layer model. Further, the head diameter from Guenkawa [7] was used, and the cylinder length was increased to ensure that the head volumes in both models are the same. For the limbs, the hand and arm volumes from the multi-layer model were combined and then used to determine the arm length in the uniform model. The same process was done for the leg in the uniform model. These anatomical characteristics, calculations, and assumptions are also discussed in more depth below.

**Table 1.** The fetal skin convective heat transfer coefficients.

| | Multi-layer cylinder model | | | | | Uniform cylinder model | | |
|---|---|---|---|---|---|---|---|---|
| Segment | L (mm) | $R_{co}$ (mm) | $R_{ms}$ (mm) | $R_{fa}$ (mm) | $R_{sk}$ (mm) | L (mm) | R (mm) | $h_i$ (W/m$^2$K) |
| Head | - | 41.0 | 46.8 | 48.3 | 50.1 | 61.0 | 53.4 | 306.8 (sphere) 224.2 (cylinder) |
| Torso | 190 | 43.2 | 46.4 | 53.4 | 54.0 | 190 | 54.0 | 222.8 |
| Arm | 125 | 4.6 | 10.0 | 14.5 | 16.0 | 147 | 16.0 | 426.7 |
| Hand | 63.0 | 5.3 | 6.5 | 7.5 | 9.5 | - | - | 566.9 |
| Leg | 180 | 3.5 | 17.7 | 19.5 | 21.0 | 203 | 21.0 | 369.0 |
| Foot | 78.0 | 3.0 | 7.89 | 9.89 | 11.4 | - | - | 511.4 |

#### S.2.2 Heat transfer between the amniotic fluid, uterine wall, and maternal core

The heat transfer from the amniotic fluid to the maternal circulation encounters three resistances in series: convection in the amniotic fluid adjacent to the uterine wall $(1/h_{af})$, conduction through the avascular amnion and chorion layers $(1/h_{mem})$, and removal by perfusing myometrial maternal blood $(1/h_{perf})$. Each resistance is derived from independent physiological data and combined as the inverse of the total equivalent uterine wall convective heat transfer coefficient ($1/h_u$):

$$\frac{1}{h_u} = \frac{1}{h_{af}} + \frac{1}{h_{mem}} + \frac{1}{h_{perf}} \quad (S13)$$

Amniotic fluid convection was determined using the properties of water at 37°C. The thermal resistance of the myometrium is determined from the thermal conductivity and thickness of the myometrium wall. Maternal side resistance is determined through the mass flow rate of the uterine blood within the myometrium. The inner surface area of the uterine wall at full term ranges from 0.1 $m^2$ to 0.2 $m^2$ [14], so we use an inner uterine wall surface area of 0.15 $m^2$. Below, we detail the calculations underlying each term in the $h_u$, which ultimately yields the total convective heat transfer coefficient of 22 $W/m^2K$.

#### S.2.2.1 Convection between the uterine wall and the amniotic fluid convection

Amniotic fluid motion is driven by fetal movements rather than organized bulk flow. The fluid undergoes gentle oscillatory rocking with cavity-scale velocities substantially lower than local limb-tip velocities. Kinematic studies report fetal lower-limb angular velocities of 30–60°/s [15] with late-gestation limb lengths of 5–10 cm [16], implying local limb-tip velocities of 3–10 cm/s and a conservative cavity-scale velocity of 1 cm/s. The fluid-filled gap between the fetal surface and the uterine wall is modeled as a parallel-plate channel of width $H = 10$ mm and hydraulic diameter $D_h = 2H = 0.020$ m. For fully developed laminar flow with both boundaries at uniform temperature, the Nusselt number is 7.6 [10], yielding:

$$h_{af} = \frac{Nu \cdot k_{af}}{D_h} = 226 \ W/m^2K \quad (S14)$$

where $k_{af} = 0.6 \ W/mK$ is the thermal conductivity of amniotic fluid (approximated as that of water at 37 °C).

#### S.2.2.2 Conduction through amnion and chorion layers

The amnion and chorion together constitute an entirely avascular passive barrier of total estimated thickness $\delta = 0.5$ mm (based on amnion 70–180 μm, chorion 20–200 μm [17]) with estimated thermal conductivity $k_{mem} \approx 0.53 \ W/mK$ [18]:

$$h_{mem} = \frac{k_{mem}}{\delta} = \frac{0.53}{5 \times 10^{-4}} = 1060 \ W/m^2K \quad (S15)$$

#### S.2.2.3 The maternal side "heat sink" limited by the myometrial blood perfusion

Prior to labor, the total uterine blood flow is 500–750 mL/min [19], of which the non-placental myometrium receives approximately 10%. The resulting wall perfusion flow is $\dot{V}_{wall} = 50$–75 mL/min. Therefore, the perfusion-limited equivalent heat transfer coefficient is:

$$h_{perf} = \frac{\dot{m} \, c_p}{A_{uterus}} = \frac{\rho_b \, \dot{V}_{wall} \, c_p}{A_{uterus}} \quad (S16)$$

Using $\rho_b = 1050 \ kg \, m^{-3}$, $c_p = 3617 \ J \, kg^{-1} K^{-1}$, $A_{uterus} = 0.15 \ m^2$, and the midpoint flow $\dot{V}_{wall} = 62.5$ mL/min gives $h_{perf} = 26.5 \ W/m^2K$ range: ~21–32 $W/m^2K$). The three resistances in series yield $h_u$ in the range of 19 to 27 $W/m^2K$, dominated by myometrial perfusion (87% of total resistance); we adopt $h_u = 22 \ W/m^2K$.

### S.3 Formulation and implementation of the two fetal thermoregulation models

#### S.3.1 Uniform cylinder model formulation and implementation

Guenkawa et al. [7] modeled the human fetus as six uniform cylinders, illustrated in Figure 1b of the main paper (1-head, 2-torso, 3-4 arms, and 5-6 legs), each exchanging blood with a central blood pool representing the heart, lungs, and liver. Under the assumption of negligible heat transfer between the umbilical cord and the amniotic fluid, we can solve for the two main unknowns, the central blood pool temperature ($T_a$) and the amniotic fluid temperature ($T_{af}$), from the following heat rate balances on the central blood pool and on the amniotic fluid [7]:

$$\sum_{i=1}^{6} m_i c_b (T_a - T_{v,i}) + m_{UA} c_b T_a - m_{UV} c_b T_{UVout} = 0 \tag{S17}$$

$$\sum_{i=1}^{6} h_i A_{s,i} (T_{s,i} - T_{af}) + h_u A_u (T_m - T_{af}) = 0 \tag{S18}$$

With the maternal core temperature being known, the above two equations can be solved once the segmental surface ($T_{s,i}$), segmental venous blood ($T_{v,i}$), and umbilical cord venous outlet ($T_{UVout}$ , which is the temperature of the blood returning to the fetus from the placenta via the cord) temperatures are expressed as functions of $T_a$ and $T_{af}$. Next, we describe solutions for the temperature distribution within each uniform cylinder to obtain $T_{s,i}$ and for the heat rate balance of each cylinder to obtain $T_{v,i}$.

The radial direction (r) steady-state bioheat transfer equation for each homogeneous cylinder with uniform volumetric heat generation ($\dot{q}_i$) exposed at $r = R_i$ to external convection with amniotic fluid (heat transfer coefficient $h_i$ and fluid temperature $T_{af}$) is [10]:

$$k_i \left(\frac{d^2 T_i}{dr^2} + \frac{dT_i}{r dr}\right) + \omega_i \rho_b c_b (T_a - T_i(r)) + \dot{q}_i = 0 \tag{S19}$$

$$\frac{dT_i}{dr}\Big|_{r=0} = 0, \quad -k_i \frac{dT_i}{dr}\Big|_{r=R_i} = h_i (T_{s,i} - T_{af}) \tag{S20}$$

Where $k_i$ is the tissue thermal conductivity (W/mK), $T_i(r)$ is the temperature at a distance r from the axis of the ith element, $\omega_i$ is the volumetric tissue blood perfusion rate (1/s), $\rho_b$ is the blood density (kg/m$^3$), $c_b$ is the blood specific heat (J/kgK), and $T_a$ is the arterial blood temperature assumed to be equal to that of the central blood pool. The closed-form solution for the two above Equations is [20]:

$$T_i(r) = T_{af} + \left(\frac{\dot{q}_i}{\omega_i \rho_b c_b} + T_a - T_{af}\right)\left(1 - \frac{h_i I_0(c_i r)}{k_i c_i I_1(c_i R_i) + h_i I_0(c_i R_i)}\right) \tag{S21}$$

Where $I_0$ and $I_1$ are zero and first-order modified Bessel functions of the first kind and:

$$c_i = \sqrt{\frac{\omega_i \rho_b c_b}{k_i}} \tag{S22}$$

To obtain the surface temperatures of each element, $T_{s,i}(R_i)$, we substitute $r = R_i$, which leads to:

$$T_{s,i} = T_{af} + \left(\frac{\dot{q}_i}{\omega_i \rho_b c_b} + T_a - T_{af}\right)\left(1 - \frac{h_i I_0(c_i R_i)}{k_i c_i I_1(c_i R_i) + h_i I_0(c_i R_i)}\right)$$

or

$$T_{s,i} = T_a Z_{1,i} + T_{af}(1 - Z_{1,i}) + Z_{2,i} \tag{S23}$$

Where

$$Z_{1,i} = \left(1 - \frac{h_i I_0(c_i R_i)}{k_i c_i I_1(c_i R_i) + h_i I_0(c_i R_i)}\right) \tag{S24}$$

and

$$Z_{2,i} = \frac{\dot{q}_i}{\omega_i \rho_b c_b} Z_{1,i} \tag{S25}$$

To solve for $T_{v,i}$, Guenkawa et al. [7] performed a heat rate balance on each body segment with surface area $A_{s,i}$:

$$\dot{q}_i V_{s,i} = h_i A_{s,i}\left(T_{s,i} - T_{af}\right) + V_{s,i}\omega_i \rho_b c_b \left(T_a - T_{v,i}\right) \tag{S26}$$

Which can be divided by the segment volume ($V_{s,i}$) and rearranged to obtain the amniotic fluid to venous temperature difference:

$$T_a - T_{v,i} = \frac{2h_i}{\omega_i \rho_b c_b R_i}\left(T_{s,i} - T_{af}\right) - \frac{\dot{q}_i}{\omega_i \rho_b c_b} \tag{S27}$$

The above equation can be expressed only as a function of $T_a$ and $T_{af}$ once the solution for $T_{s,i}$ is substituted:

$$T_a - T_{v,i} = T_a Z_{1,i} Z_{3,i} + T_{af}\left(-Z_{1,i} Z_{3,i}\right) + Z_{4,i} \tag{S28}$$

Where

$$Z_{3,i} = \frac{2h_i}{\omega_i \rho_b c_b R_i} \tag{S29}$$

and

$$Z_{4,i} = Z_{2,i} Z_{3,i} - \frac{\dot{q}_i}{\omega_i \rho_b c_b} \tag{S30}$$

Substituting Eqs. S8, S23, and S27 into the amniotic fluid and central blood pool heat rate balances (Eqs. S17 and S18) and manipulating the equations to isolate the temperatures, we obtain:

$$T_a w_{11} + T_{af} w_{12} + M_1 = 0 \tag{S31}$$

$$T_a w_{21} + T_{af} w_{22} + M_2 = 0 \tag{S32}$$

Where

$$w_{11} = \sum_{i=1}^{6} h_i A_{s,i} Z_{1,i} = \sum_{i=1}^{6} h_i A_{s,i}\left(1 - \frac{h_i I_0(c_i R_i)}{k_i c_i I_1(c_i R_i) + h_i I_0(c_i R_i)}\right) \tag{S33}$$

$$w_{12} = -\sum_{i=1}^{6} h_i A_{s,i} Z_{1,i} - h_u A_u = -\sum_{i=1}^{6} h_i A_{s,i}\left(1 - \frac{h_i I_0(c_i R_i)}{k_i c_i I_1(c_i R_i) + h_i I_0(c_i R_i)}\right) - h_u A_u \tag{S34}$$

$$M_1 = \sum_{i=1}^{6} h_i A_{s,i} Z_{2,i} + h_u A_u T_m = \sum_{i=1}^{6} \frac{\dot{q}_i h_i A_{s,i}}{\omega_i \rho_b c_b}\left(1 - \frac{h_i I_0(c_i R_i)}{k_i c_i I_1(c_i R_i) + h_i I_0(c_i R_i)}\right) + h_u A_u T_m \tag{S35}$$

$$w_{21} = \sum_{i=1}^{6} \omega_i \rho_b c_b \pi R_i^2 L_i\, Z_{1,i} Z_{3,i} + m_{UA} c_b - m_{UV} c_b Z_5 =$$

$$\sum_{i=1}^{6} 2 h_i \pi R_i L_i \left(1 - \frac{h_i I_0(c_i R_i)}{k_i c_i I_1(c_i R_i) + h_i I_0(c_i R_i)}\right) + m_{UA} c_b$$

$$- m_{UV} c_b \left(\varepsilon_c + \frac{\left(1-\varepsilon_p\right)(1-\varepsilon_c)^2}{1-\varepsilon_c\left(1-\varepsilon_p\right)}\right) \tag{S36}$$

$$w_{22} = -\sum_{i=1}^{6} \omega_i \rho_b c_b \pi R_i^2 L_i\, Z_{1,i} Z_{3,i} = -\sum_{i=1}^{6} 2 h_i \pi R_i L_i \left(1 - \frac{h_i I_0(c_i R_i)}{k_i c_i I_1(c_i R_i) + h_i I_0(c_i R_i)}\right) \tag{S37}$$

$$M_2 = \sum_{i=1}^{6} \omega_i \rho_b c_b \pi R_i^2 L_i\, Z_{4,i} - m_{UA} c_b Z_6 =$$

$$\sum_{i=1}^{6} \pi \dot{q}_i R_i^2 L_i \left(\frac{2 h_i}{\omega_i \rho_b c_b R_i}\left(1 - \frac{h_i I_0(c_i R_i)}{k_i c_i I_1(c_i R_i) + h_i I_0(c_i R_i)}\right) - 1\right)$$

$$- m_{UA} c_b \left(\frac{\varepsilon_p (1-\varepsilon_c)}{1-\varepsilon_c\left(1-\varepsilon_p\right)}\right) \tag{S38}$$

and

$$Z_{4,i} = \left(1 - \frac{h_i I_0(c_i R_i)}{k_i c_i I_1(c_i R_i) + h_i I_0(c_i R_i)}\right) \frac{\dot{q}_i}{\omega_i \rho_b c_b} \frac{2 h_i}{\omega_i \rho_b c_b R_i} - \frac{\dot{q}_i}{\omega_i \rho_b c_b} \tag{S39}$$

$$\frac{\dot{q}_i}{\omega_i \rho_b c_b} \left(\left(1 - \frac{h_i I_0(c_i R_i)}{k_i c_i I_1(c_i R_i) + h_i I_0(c_i R_i)}\right) \frac{2 h_i}{\omega_i \rho_b c_b R_i} - 1\right) \tag{S40}$$

$$M_2 = \sum_{i=1}^{6} \omega_i \rho_b c_b \pi R_i^2 L_i\, Z_{4,i} - m_{UA} c_b Z_6 =$$

$$\sum_{i=1}^{6} \pi \dot{q}_i R_i^2 L_i \left(\frac{2 h_i}{\omega_i \rho_b c_b}\left(1 - \frac{h_i I_0(c_i R_i)}{k_i c_i I_1(c_i R_i) + h_i I_0(c_i R_i)}\right) - 1\right)$$

$$- m_{UA} c_b \left(\frac{\varepsilon_p (1-\varepsilon_c)}{1-\varepsilon_c\left(1-\varepsilon_p\right)}\right) \tag{S41}$$

The above equations can be readily solved for $T_a$ and $T_{af}$ as:

$$T_a = \frac{\frac{M_2}{w_{22}} - M_1}{w_{11} - \frac{w_{21}}{w_{22}}} \tag{S42}$$

$$T_{af} = \frac{-T_a w_{21} - M_2}{w_{22}} \tag{S43}$$

**S.3.2 Multi-compartmental cylinder model formulation and implementation**

The fetal nodes have basal metabolic heat generation, heat transfer by blood perfusion,conduction between the tissue layers,and convective heat transfer between the skin and amniotic fluid. With each tissue layer treated as a single lumped thermal node governed by a discrete form of the Pennes bioheat equation [21], the metabolic heat generation, inter-layer conduction, and convective exchange with the arterial blood supply are balanced at steady state:

$$\underbrace{\dot{q}_{j,i}\, V_{lay(i,j)}}_{\substack{\text{Basal metabolic} \\ \text{heat generation}}} + \underbrace{\rho_b\, c_b\, \omega_{j,i}\, V_{lay(i,j)} \left(T_a - T_{j,i}\right)}_{\substack{\text{Convective heat exchange between} \\ \text{tissue layer and blood}}} + \underbrace{TC_{in,j,i} \left(T_{in,j,i} - T_{j,i}\right)}_{\text{conduction heat gain}}$$

$$- \underbrace{TC_{out,j,i} \left(T_{j,i} - T_{out,j,i}\right)}_{\text{conduction heat loss}} + \underbrace{h_i\, A_{s,i} \left(T_{sk,i} - T_{af}\right)}_{\substack{\text{convection from fetal skin to} \\ \text{amniotic fluid}}} = 0 \tag{S44}$$

Where, i is body segment index (i= 1:head, 2:torso, 3:arms, 4:hands, 5:legs, 6:feet), j is tissue layer index (j= core, muscle, fat, skin), $\dot{q}_{j,i}$ is volumetric metabolic heat generation rate in layer j of segment i [W/m³], $V_{lay(i,j)}$ is volume of layer j in segment i [m³], $\rho_b$: blood density=1050 [kg/m³], $c_b$ is blood specific heat = 3617 [J/kgK], $\omega_{j,i}$ is blood perfusion rate of layer j in segment i [1/s], $T_{j,i}$ is temperature of layer j in segment i [°C], $TC_{in,j,i}$ is thermal conductance between layer j and its inward-adjacent layer [W/K] ($TC_{in,j,i} = 0$ when j= core), $T_{in,j,i}$ is temperature of the inward-adjacent layer to node $(j, i)$[°C], $TC_{out,j,i}$ is thermal conductance between layer j and its outward-adjacent layer [$WK^{-1}$] ($TC_{out,j,i} = 0$ when j= sk), $T_{out,j,i}$ is temperature of the outward-adjacent layer to node $(j, i)$ [°C]. The energy balances for each tissue layer within each segment are:

Energy balances for each layer (core, muscle, fat, and skin) and body segment(head, torso, arm, leg, hand, and foot that for the limbs these values are multiplied by two in the model once the relevant equations are solved) produce 24 equations that with adjusted versions of Eq.17-18 (for number of nodes) produce 26 linear equations and 26 unknown temperatures. These equations form a system of linear equations that are solved by matrix inversion (in matrix form: AT = b, where A is the thermal conductance matrix, T is the vector of unknown temperatures, and b is the vector of prescribed heat source and boundary condition terms).

**S.4 From fetal anatomical to cylinder (and sphere) fetal geometry**

All geometrically simplified segment dimensions correspond to a healthy fetus at 39 weeks of gestational age with a body mass of approximately 3.2 kg, representing the 50$^{th}$ percentile. We derived the dimensions from primary clinical imaging studies and term neonatal anthropometric surveys; no parameters were taken from adult anatomy unless explicitly stated. The values are provided in Table 1 above while the details of calculations and assumptions behind each segment are described below.

#### S.4.1 The head geometry

The fetal head at term is a spheroid rather than a sphere or cylinder: the biparietal diameter (BPD), occipitofrontal diameter (OFD), and suboccipital-bregmatic height differ from one another by no more than 15%. Because this aspect ratio is far closer to spherical than to cylindrical, the head was modeled as a sphere with a volume-equivalent radius:

$$R_{eff} = (abc)^{\frac{1}{3}} = \left(\frac{BPD}{2} * \frac{OFD}{2} * \frac{H}{2}\right)^{\frac{1}{3}} \quad (S45)$$

Where a, b, c are the three principal semi-axes. Using BPD = 93 mm and OFD = 113.5 mm at the 50th percentile [22][23], and H = 95 mm from the existing fetal biometry literature [24], this yields $R_{eff}$ = 50.05 mm. The volume preservation property of the geometric mean ensures that the spherical model contains the same internal volume as the true spheroid. Four concentric layers are defined within the spherical head, with thicknesses derived as follows. The scalp layer (1.8 mm) corresponds to the midpoint of the 1–2 mm range reported for fetal scalp skin at term [25]. The cranial bone layer (1.5 mm) reflects direct measurements of parietal bone thickness from neonatal cadaveric specimens and CT-based statistical models, which consistently report values of 1.3–1.5 mm at birth [26]. The core layer radius is derived from the volume of the cortical gray matter, which is 140 $cm^3$ [27]. The thickness of the muscle layer is assumed to be small for the head (1.5 mm). The radius of the core layer in the head is derived using the radius of the muscle layer and the volume of the cortical gray matter, the core radius is found utilizing:

$$r_{head,core} = \left(r_{mu}^3 - \frac{3 * V_{CGM}}{4\pi}\right)^{1/3} \quad (S46)$$

#### S.4.2 The trunk geometry

The trunk was modeled as a circular cylinder, with the outer radius determined by the abdominal circumference. The length of the torso is 190 mm [28]. The abdominal circumference is 340 mm [29]; this is converted to a radius to represent the skin layer of the torso cylindrical segment.

$$R_{sk} = \frac{circumference}{2\pi} = 54.11 \quad (S47)$$

The thickness of the skin (dermis) on the back and abdomen is approximately 0.72 mm [30], as determined by ultrasound study. The thickness of the fat layer is approximately 7 mm [31]. The thickness of the abdominal muscle layer is found by adding the three measured [32] wall muscle values: external oblique (EO) 1.02 mm (± 0.33), internal oblique (IO) 1.16 mm (± 0.39), and transversus abdominis (TA) 1.02 mm (± 0.37).

$$t_{muscle} = EO + IO + TA \quad (S48)$$

The final value for the abdominal muscle layer thickness is 3.2 mm.

#### S.4.3 The limb geometries

The arm length of 124.9 mm was determined by adding the length of the humerus and ulna, which are 63.7 mm and 61.2 mm, respectively [33]. The mid-arm circumference [34], which is approximately 100 mm, was converted to the radius of the skin layer on the arm by:

$$r_{arm,skin} = \frac{circumference_{mid\ arm}}{2\pi} = 16\ mm \quad (S49)$$

The core of the arm is determined by taking the diameter of the ulna and the radius and adding them together the diameter of the radius is 5.6 mm [35] and the diameter of the ulna is 3.6 mm

[36]so the resulting radius of the core of the arm is 4.6 mm. Arm fat thickness is measured using the skinfold thickness and dermis thickness. The skinfold of the triceps is estimated to be between 4-5 mm [37,38]; therefore, we use the average thickness of 4.5 mm. Utilizing the triceps skinfold thickness of 4.5 mm and dermis thickness of 2 mm, the arm fat thickness is calculated as:

$$t_{arm,fat} = \frac{tricep_{skinfold} + t_{dermis}}{2} \quad (S50)$$

The skin thickness for the arm is 1.5 mm [39]. The muscle thickness is determined by difference:

$$t_{mu} = R_{sk} - R_{co} - t_{fa} - t_{sk} = 5.4 \text{ mm} \quad (S51)$$

Malas et al. found the average length of a fetal hand (from wrist to middle finger) is 63 mm [40]. The radius for the hand cylindrical body segment is found by taking the geometric mean of the hand width and skin depth at the metacarpal level. The thickness of the fat layer on the hand is assumed to be less than that of the arm and is estimated at 1 mm because there is very little subcutaneous fat on a fetus hand [41]. The skin layer thickness is 2 mm [42]. the thickness of muscle was determined by taking the ratio of muscle thickness of arm muscle thickness on an adult arm compared to the muscle thickness on the adult hand [43], this percentage was found to be about 22% so taking the fetal muscle thickness for the arm and multiplying it by the ratio found from adult physiology, the muscle thickness of the hand is estimated to be 1.2 mm. The width of a hand (distance between the outer joints of the 2$^{nd}$ and 5$^{th}$ metacarpophalangeal (knuckles)) is 38 mm [40], because this value is just of the bones we also need the skin depth at the metacarpal level which is found by adding the thicknesses of the muscle, fat, and skin on the hand. The thickness of the bone was then found by dividing the length of the metacarpophalangeal by four to account for each finger, then this value accounted for the thickness of one finger and when the thickness of the fat, muscle, and skin was subtracted from this value the core thickness was found, which is 5.3 mm.

To estimate the outer radius of the leg cylindrical segment, the average of the thigh (155.5 mm) [44] and calf (108.8 mm) [34] circumferences are transformed into a radius by:

$$r_{leg,skin} = \frac{(circumference_{thigh} + circumference_{calf})/2}{2\pi} \quad (S52)$$

We set the length of the fetal leg at 180 mm [28] and its core at 7 mm, which is minimum diaphysis diameter for a newborn [45]. Following the same approach used for the arms, the thickness of the leg fat layer was determined from skinfolds and the dermal thicknesses. The skinfold thickness of the legs is 5 mm [46], so the thickness of the fat layer was set to 1.8 mm while that of the skin layer was set to 1.5 mm.

The length of a fetal foot is 78.2 mm [33]. In particular, we utilized an equation for the full foot width (FFW) for a fetus from [47]. The width of the foot is 20.76 mm, and the dorsoplantar depth is assumed to be double the size of the hand depth at the metacarpal level (20 mm). We used a geometric mean of these values to find the outer radius of the foot cylinder:

$$r_{foot,sk} = \sqrt{\frac{width}{2} + \frac{depth}{2}} = 11.4 \text{ mm} \quad (S53)$$

We set the radius of the core layer in the foot based on the diameter of the talus bone which is 6 mm [48] so the radius of the core layer will be 3 mm. We set the foot skin layer thickness to 1.5 mm, and its fat layer thickness to 2 mm.

### S.5 The blood flow rate and distribution within the fetus

The near-term fetal combined cardiac output (CCO) can be described using the double-exponential relationship [4]:

$$\mathrm{CCO(t)} = 3400.88(1.141 * 10^{-5})^{e^{(0.07022(t-2))}} = 1544\frac{\mathrm{mL}}{\mathrm{min}} @t = 39 \text{ weeks} \tag{S54}$$

The fraction of this total blood flow going into the brain and the placenta can be estimated using similar empirical fits [4], for example providing a placental CCO fraction near 16% at term. However, since many Doppler and MRI studies [49,50] report umbilical/placental flow closer to 20–22% of CCO near term, we use a rounded value of 20%. With a total CCO of 1544 mL/min, the placental flow is 309 mL/min. Similarly, we can estimate that the brain receives about 18% of the CCO; however, determining fractions for other body segments is harder, since measurements typically focus on specific organs (e.g., lungs, liver, intestine, and kidney).

We estimated the remaining segmental blood flow from aortic flow measurements. The ascending aortic flow is 41% CCO while the descending aortic flow is 55% CCO at term [4]. Because the descending aorta supplies both the placenta, the liver, and the lower body, subtracting placental and hepatic flows from descending aortic flow gives approximately 27% remaining for the fetal lower body itself. This lower-body flow must perfuse the abdominal organs, pelvis, and lower extremities, implying that the legs likely receive only part of this amount, approximately 15–20% CCO (230–310 mL/min). Similarly, the ascending aorta supplies the brain, myocardium, upper torso, and upper extremities. Since carotid flow alone is approximately 21% CCO and coronary flow is roughly 3–5% CCO, only a modest residual fraction remains available for the arms, yielding an estimated upper-extremity perfusion of roughly 3–7% CCO (45–110 mL/min).

In summary, for both models, the approximate flow percentages for each body segment are as follows: placenta 20%, head 22%, torso 34%, each arm-3%, each leg 9%, with further breakdown summarized in Tables 2 and 3 below.

**Table 2.** Thermophysical and physiological tissue parameters for uniform cylinder model.

| Segment | k | $\dot{q}$ | Q (total 8.04W) | blood flow (total 1544 ml/min) | ω |
|---|---|---|---|---|---|
| | (W/mK) | (W/m³) | (W) | (% CCO) | (1/s) |
| Head | 0.53 | 8474.2 | 4.63 | 21.7 | 0.01 |
| Torso | 0.43 | 1700.9 | 2.96 | 33.7 | 0.005 |
| Arm | 0.37 | 551.6 | 0.065 | 3 | 0.0065 |
| Arm | 0.37 | 551.6 | 0.065 | 3 | 0.0065 |
| Leg | 0.38 | 562.6 | 0.16 | 9 | 0.0082 |
| Leg | 0.38 | 562.6 | 0.16 | 9 | 0.0082 |

For the multi-layer model, flow is further partitioned among the four tissue layers according to physiologically guided layer fractions (Table 3); the layer-specific perfusion rate $\omega_{ij}$[1/s] is then computed as:

$$\omega_{ij} = \frac{f_{ij}\,\dot{V}_{seg,i}}{V_{ij}} \tag{S55}$$

where $f_{il}$is the layer flow fraction, $\dot{V}_{seg,i}$the bilateral segment flow, and $V_{ij}$the bilateral layer volume. The resulting perfusion conductance entering the matrix are:

$$\Phi_{ij} = \omega_{ij}\,\rho_b\,c_b\,V_{ij}[\mathrm{W/K}] \tag{S56}$$

**Table 3.** Thermophysical and physiological tissue parameters for multi-layer model [18,51–55] (BL=blood flow).

| Segment / Layer | | k | $\dot{q}$ | TC | BL | Layer flow fraction | | | |
|---|---|---|---|---|---|---|---|---|---|
| | | (W/mK) | (W/m$^3$) | (W/K) | (%CCO) | $f_{co}$ (%) | $f_{ms}$ (%) | $f_{fa}$ (%) | $f_{sk}$ (%) |
| Head | Core | 0.565 | 15,553 | 0.30 | 21.7 | 80 | 12 | 4 | 4 |
| | Muscle | 0.565 | 686 | 7.72 | | | | | |
| | Fat | 0.210 | 199 | 5.79 | | | | | |
| | Skin | 0.370 | 644 | - | | | | | |
| Torso | Core | 0.510 | 2,547 | 8.11 | 33.7 | 90 | 4 | 3 | 3 |
| | Muscle | 0.490 | 686 | 2.57 | | | | | |
| | Fat | 0.210 | 199 | 7.05 | | | | | |
| | Skin | 0.370 | 644 | - | | | | | |
| Arms† | Core | 0.320 | 394 | 0.54 | 4.76 | 5 | 85 | 5 | 5 |
| | Muscle | 0.490 | 686 | 3.45 | | | | | |
| | Fat | 0.210 | 199 | 4.10 | | | | | |
| | Skin | 0.370 | 644 | - | | | | | |
| Hands† | Core | 0.320 | 394 | 0.34 | 0.94 | 5 | 80 | 8 | 7 |
| | Muscle | 0.490 | 686 | 2.04 | | | | | |
| | Fat | 0.210 | 199 | 2.12 | | | | | |
| | Skin | 0.370 | 644 | - | | | | | |
| Legs† | Core | 0.320 | 394 | 0.54 | 15.67 | 5 | 85 | 5 | 5 |
| | Muscle | 0.490 | 686 | 0.60 | | | | | |
| | Fat | 0.210 | 199 | 0.94 | | | | | |
| | Skin | 0.370 | 644 | - | | | | | |
| Feet† | Core | 0.320 | 394 | 0.79 | 2.03 | 5 | 75 | 10 | 10 |
| | Muscle | 0.490 | 686 | 0.64 | | | | | |
| | Fat | 0.210 | 199 | 0.93 | | | | | |
| | Skin | 0.370 | 644 | - | | | | | |

† Bilateral segment;

### S.6 Thermal conductivity and inter-layer conductance of the fetal tissues

The thermal conductivities for each tissue type were obtained from the IT'IS database of thermal properties for biological tissues [18], with tissue-volume-weighted averages used for the uniform cylinder model. Within the multi-layer model, heat transfer between tissue layers is determined using the thermal conductance (TC), which also accounts for the thicknesses of the tissue layers. In particular, the heat transfer between layers is calculated as:

$$h_{cond_{i,i'}} = TC_{i,i'}(T_i - T_{i'}) \tag{S57}$$

The thermal conductance is determined by using the tissue thermal conductivities ($k_i$), which are used to calculate the effective conductivity for the interface between the tissue layers ($k_{eff}$) using:

$$k_{eff} = \frac{2k_i k_{i'}}{k_i + k_{i'}} \tag{S58}$$

In turn, the radial thermal conductance between the layers is calculated using:

$$TC_{cylinder} = \frac{2\pi k_{eff} L}{\ln\left(\frac{r_{outer}}{r_{inner}}\right)} \tag{S59}$$

$$TC_{sphere} = \frac{4\pi k_{eff} r_{outer} r_{inner}}{r_{outer} - r_{inner}} \tag{S60}$$

For the thermal conductance interface between the core and muscle, the inner radius is zero, so the equations for these layers utilize the radius of the core:

$$TC_{head,cm} = 4\pi r_{core} \tag{S61}$$

$$TC_{cyl,cm} = 2\pi k_{eff} L \tag{S62}$$

**Table 4.** Thermal conductivities and conductance for each layer and body Segment [18].

| Segment | Layer | $k_i$ [W/mK] | TC [W/K] |
|---|---|---|---|
| **Head** | Core | 0.565 | 0.30 |
| | Muscle | 0.565 | 7.72 |
| | Fat | 0.320 | 5.79 |
| | Skin | 0.370 | - |
| **Torso** | Core | 0.510 | 8.11 |
| | Muscle | 0.490 | 2.57 |
| | Fat | 0.210 | 7.05 |
| | Skin | 0.370 | - |
| **Leg**$^+$ | Core | 0.400 | 0.54 |
| | Muscle | 0.490 | 3.45 |
| | Fat | 0.210 | 4.10 |
| | Skin | 0.370 | - |
| **Arm**$^+$ | Core | 0.400 | 0.34 |
| | Muscle | 0.510 | 2.04 |
| | Fat | 0.490 | 2.12 |
| | Skin | 0.210 | - |
| **Hand**$^+$ | Core | 0.400 | 0.54 |
| | Muscle | 0.490 | 0.60 |
| | Fat | 0.210 | 0.94 |
| | Skin | 0.370 | - |
| **Foot**$^+$ | Core | 0.400 | 0.79 |
| | Muscle | 0.490 | 0.64 |
| | Fat | 0.210 | 0.93 |
| | Skin | 0.370 | - |

+: TC represents the value of one limb

#### S.7 Metabolic heat generation of the fetal tissues

Elia et al. [51] estimated the metabolic rate of each organ and tissue within the body. For the muscle, fat, and skin layers, the specific metabolic rate is the same for each segment. For the core layer, the segments have varying specific metabolic rates due to differences in the bone-to-viscera ratio and the viscera within each body segment. Torso core (visceral organ mix):

$$Q_{organs} = \sum K_i * m_i \tag{S63}$$

Organ masses were extrapolated from Table **5** in Gruenwald et al. [56]. The metabolic rate for each organ was found using Table 1 in the paper by Elia [51].

**Table 5.** The mass and specific metabolic rate of tissues within the torso core used to determine effective metabolic rate for the segment.

| Organ | Mass [g] | Specific Metabolic Rate [kcal/kg/day] |
|---|---|---|
| **Liver** | 141.4 | 200 |
| **Heart** | 21.34 | 440 |
| **Kidney** | 26.78 | 440 |
| **Lung** | 57.8 | 12 |
| **Spleen** | 10.82 | 12 |

Converting from kcal/kg/day to W/kg:

$$\frac{1\,\mathrm{kcal/kg}}{\mathrm{day}} * 4184\,\mathrm{J/kcal} \div 86400\,\mathrm{s/day} = 0.0483\,\mathrm{W/kg} \tag{S64}$$

$$\mathrm{Q_{organs}} = (200 * 0.1414 + 440 * 0.02134 + 440 * 0.02678 + 12 * 0.0578 + 12 * 0.01082) * 0.04843 = 2.435\,\mathrm{W} \tag{S65}$$

$$\mathrm{V_{core,torso}} = 1048\,\mathrm{cm^3} \tag{S66}$$

$$\dot{\mathrm{q}}_{\mathrm{core,torso}} = \frac{\mathrm{Q_{organs}}}{\mathrm{V_{core,torso}} * 10^{-6}} = 1{,}645\,\mathrm{W/m^3} \tag{S67}$$

To determine the metabolic heat generation within the limb cores (Arms, Hands, Legs, Feet), with assumed that bone composition is 60% cortical and 40% marrow and utilize the below equations:

$$\dot{\mathrm{q}}_{\mathrm{cortical}} = 3\frac{\mathrm{kcal/kg}}{\mathrm{day}} * 1908\,\mathrm{kg/m^3} * 0.04843 = 277\,\mathrm{W/m^3} \tag{S68}$$

$$\dot{\mathrm{q}}_{\mathrm{marrow}} = 12\frac{\mathrm{kcal/kg}}{\mathrm{day}} * 980\,\mathrm{kg/m^3} * 0.04843 = 569\,\mathrm{W/m^3} \tag{S69}$$

$$\dot{\mathrm{q}}_{\mathrm{core,limbs}} = 0.6 * 277 + 0.4 * 569 = 394\ \mathrm{W/m^3} \tag{S70}$$

For muscle, fat, and skin, the metabolic rates are, 13, 4.5, and 12 $\frac{\mathrm{kcal/kg}}{\mathrm{day}}$, respectively [51]. The metabolic heat generation values for muscle, fat, and skin are then found to be, 686, 199, and 644 $\mathrm{W/m^3}$, respectively.

### S.8 Radial temperature variation in hands, legs, and feet

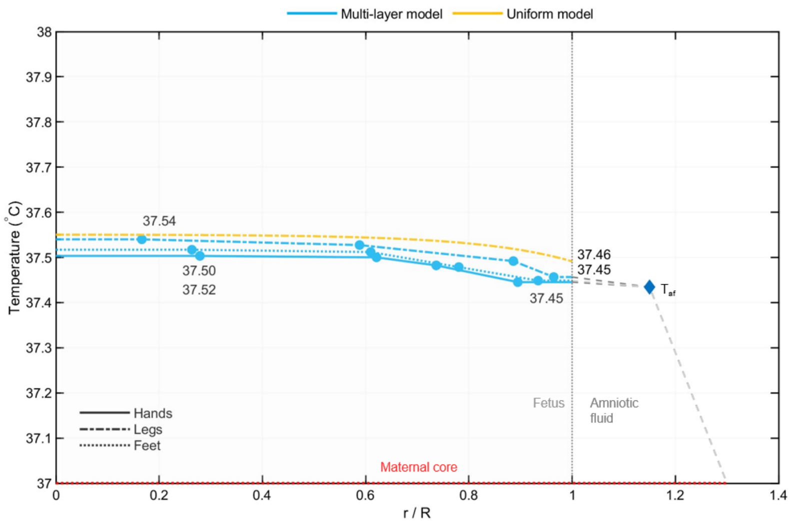


**Figure 1.** predicted radial temperature variation in hands, legs, and feet as function of scaled radius (r/R)

### S.9 MATLAB code implementation of uniform model

```
function out = UniformModelFunction(Tmother)
%Author: Konrad Rykaczewski,
% Code implementing Guenkawa and Ferreira, Steady-state Temperature
% Distribution in a Fetus, Proceedings of 15th Intenratinoal Conference on
% Heat Transfer, Fluid Mechanics, and thermodynamics, pg.2039 to 2044, 2021
% Usage: out = UniformModelFunction(Tmother)
%returns
% Returns fetal center temperature (segment 1 center) and outputs OR radial temperature variation
Radius/50 steps in head, torso, arm, and
%leg columns


%% ---------- Physical constants &  ----------
rho_b = 1050;        % kg/m3
c_b   = 3617;        % J/kgK

mUA = 0.00556;        % umbilical arterial mass flow (kg/s)=318 mL/min or
mUV = 0.00556;        % umbilical venous mass flow (kg/s)

Ec=0.32;%umbilical cord counterflow heat exchanger effectivness

Ep=0.845; %placenta effectivness
Z5=Ec+((1-Ep)*(1-Ec)^2)/(1-Ec*(1-Ep));
Z6=(Ep*(1-Ec))/(1-Ec*(1-Ep));

h_uterus = 22;           % W/m2K uterine conv
A_uterus = 0.15;         % m2 uterine inner area

%% ---------------- Guenkawa table updated with more realistic physiological parameters(6
segments) ----------------
% Columns: [R (m), L (m), k (W/mK), omega (1/s), qdot (W/m3), h (W/m2K)]
%Rows: head,torso,arm,arm,leg,leg]
seg = [
    0.0534,  0.061, 0.53, 0.010, 8474.2, 224;
    0.054,  0.190, 0.43, 0.005,  1700.9, 222.8;
    0.016,  0.147, 0.37, 0.0065, 551.6, 427;
    0.016,  0.147, 0.37, 0.0065, 661.6, 427;
    0.021,  0.203, 0.38, 0.0082, 562.6, 369;
    0.021,  0.203, 0.38, 0.0082, 562.6, 369
];
R = seg(:,1); Lseg = seg(:,2); k = seg(:,3);
omega = seg(:,4); qdot = seg(:,5); h = seg(:,6);

% Derived geometric quantities (exact formulae)
A_surf = 2*pi .* R .* Lseg;            % segment surface area

%% ------- Bessel-based closed-form segment coefficients -------
c_i = sqrt(omega.* rho_b* c_b./k);
z = c_i .* R;
I0 = besseli(0,z);
I1 = besseli(1,z);

Z1i=1-(h.*I0)./(c_i.*k.*I1+h.*I0);
Z2i=qdot.*Z1i./(omega*rho_b*c_b);


%amniotic fluid energy balance coefficients aW11*Ta+aW12*Taf=-aC1
aW11=sum(h.*A_surf.*(1-(h.*I0)./(c_i.*k.*I1+h.*I0)));
aW12=-sum(h.*A_surf.*(1-(h.*I0)./(c_i.*k.*I1+h.*I0)))-h_uterus*A_uterus;
aC1=sum(h.*A_surf.*qdot.*(1-
(h.*I0)./(c_i.*k.*I1+h.*I0))./(omega*rho_b*c_b))+h_uterus*A_uterus*Tmother;

%blood reservoir energy balance coefficients: aW21*Ta+aW22*Taf=-aC2
aW21=sum(2*3.141.*h.*R.*Lseg.*(1-(h.*I0)./(c_i.*k.*I1+h.*I0)))+mUA*c_b-mUV*c_b*Z5;
aW22=-sum(2*3.141.*h.*R.*Lseg.*(1-(h.*I0)./(c_i.*k.*I1+h.*I0)));
aC2=sum(3.141.*qdot.*R.*R.*Lseg.*(2*h./(R.*omega*rho_b*c_b).*(1-(h.*I0)./(c_i.*k.*I1+h.*I0))-1))-
mUA*c_b*Z6*Tmother;
```

```
%directly solved
Taf=(aC1*aW21/aW11-aC2)/(aW22-aW21*aW12/aW11);
Ta=(-Taf*aW12-aC1)/aW11;

%head center segment (segment 1) center temp closed form
i = 1;
c1 = sqrt(omega(i) * rho_b * c_b / k(i));
z1 = c1 * R(i);
I0_1 = besseli(0,z1); I1_1 = besseli(1,z1);
D1 = k(i) * c1 * I1_1 + h(i) * I0_1;
T_part1 = qdot(i) / ( omega(i) * rho_b * c_b );
Cpref1 = - h(i) / D1;
T_center = (Ta + T_part1) + Cpref1 * ((Ta + T_part1) - Taf);
T_fetus = T_center;
Tshead=Ta*Z1i(i)+Taf*(1-Z1i(i))+Z2i(i);
Z1i=1-(h.*I0)./(c_i.*k.*I1+h.*I0);
Z2i=qdot.*Z1i./(omega*rho_b*c_b);

%output head center, blood pool, amniotic fluid, scalp (skin on the head) temperatures
%out=[T_fetus, Ta, Taf, Tshead];

%instead of above output, here we output the radial variation of
%radial temperature variation in Radius/50 steps in head, torso, arm, and
%leg columns
ivals=1:51;
out=zeros(length(ivals),4);
cols=[1 2 3 5];
for j=1:length(ivals)
    for ki=1:4
        i = cols(ki); % actual loop value (1,2,3,5)
        c1 = sqrt(omega(i) * rho_b * c_b/k(i));
        z1 = c1 * R(i);
        I0_1 = besseli(0,z1); I1_1 = besseli(1,z1);
        I0_r=besseli(0,c1*R(i)*(j-1)/50);
        D1 = k(i) * c1 * I1_1 + h(i) * I0_1;
        T_part1 = qdot(i) / ( omega(i) * rho_b * c_b );
        Cpref1 = - h(i)*I0_r / D1;
        T_r= (Ta + T_part1) + Cpref1 * ((Ta + T_part1) - Taf);
        out(j,ki)=T_r;% store in column 1-4
    end
end
end
```

### S.10 MATLAB code implementation of multilayer cylinder model

To properly execute the below code, put the file in the same folder as the above uniform-cylinder model (with file name adjusted to function name) and with the validation data form Lavesson et al. replicated in Section S.11 below. Executing the code replicates Figure 2a and radial temperature distributions in all body segments.

```
%% MultiLayerModelWithUniformModel_final.m
%  Steady-state fetal thermal multi-layer model w uniform — Final
%  6 segments × 4 layers + T_a + T_af  →  26×26 linear system
%
%  Segments:
%    1. Head  (sphere)
%    2. Torso (cylinder)
%    3. Arms  (bilateral pair — 2 × single arm)
%    4. Hands (bilateral pair — 2 × single hand)
%    5. Legs  (bilateral pair — 2 × single leg)
%    6. Feet  (bilateral pair — 2 × single foot)
%
%  Layer order (all segments):  1=Core · 2=Muscle · 3=Fat · 4=Skin
%
%
%  Geometry:
%    • HEAD modelled as a SPHERE instead of a cylinder.
%      The fetal head is a prolate spheroid (BPD/OFD differ by 25%), far
%      closer to spherical than cylindrical.  The equivalent sphere radius
%      R_eff = ∛(a·b·c) preserves the true ellipsoidal volume.
%      Spherical-shell TC formula replaces cylindrical-shell formula for
%      all four head layers.
%    • All segment dimensions re-derived from clinical imaging and neonatal
%      anthropometry (see §GEOMETRY for full source citations; Table 1).
%    • Arm, Hand, Leg, Foot are NEW bilateral segments replacing the four
%      separate L/R segments of v1.  Bilateral factors double V, TC, Φ, A.
%
%  Thermal conductivities (Table 2, finalized; IT'IS Hasgall et al. 2022):
%    • Head core k=0.565 (brain+skull composite), Torso core k=0.510
%      (visceral organ composite), limb cores (Arms/Hands/Legs/Feet)
%      k=0.320 (cortical+marrow bone composite), non-Head muscle/scalp
%      muscle k=0.490.  Fat k=0.210, skin k=0.370 (all segments).
%
%  Metabolic heat:
%    • qdot computed from tissue-specific Ki (Elia 1992) × density (IT'IS 2022)
%      rather than from Guenkawa calibration values.
%    • Head core (brain+skull lumped) uses whole-brain Ki (Elia 1992,
%      240 kcal/kg/day) over brain mass (Andescavage 2017) → qdot_co=15,553 W/m³.
%    • Torso core uses organ-weighted Q from Gruenwald (1966) masses
%      → qdot_co=2547 W/m³ with the finalized Table 1 core volume.
%    • Limb cores (60% cortical/295 + 40% marrow/455 W/m³, Hasgall et al.
%      2022) → qdot_co_limb=359 W/m³.
%    • Fat and skin qdot corrected from Fiala values to Elia/IT'IS values
%      (fat: 58→199 W/m³; skin: 368→644 W/m³).
%
%  Combined ventricular output (CVO):
%    • CVO partition updated from Rudolph (1985, ovine, 450 mL/min/kg,
%      22/34/6/18%) to Mielke & Benda (2001, human Doppler, 425 mL/min/kg,
%      21.7/33.7/5.7/17.7%).
%
%  Boundary conditions:
%    • Uterine wall:  h_uterus = 22 W/m²K, A_uterus = 0.15 m²
%    • Umbilical flow: mUA = 5.62×10⁻³ kg/s
%      318 mL/min (Zhang and Lindsey)
%  ──────────────────────────────────────────────────────────────────────

clearvars; clc; close all;

out_dir = 'figures_v3';
if ~exist(out_dir,'dir'), mkdir(out_dir); end
```

```
%% ========================================================================
%  PHYSICAL CONSTANTS
%  ρ_blood: IT'IS Foundation (Hasgall et al. 2022), human blood at 37 °C
%  c_blood: Poppendiek et al. (1966) — commonly used in Pennes-BHE models
%% ========================================================================

rho_b = 1050;      % blood density           [kg/m³]
c_b   = 3617;      % blood specific heat     [J/kg·K]

%% ========================================================================
%  MATERNAL CORE TEMPERATURE
%% ========================================================================

Tmother = 37.0;    % maternal core temperature  [°C]

%% ========================================================================
%  SEGMENT GEOMETRY  (6 segments, 4 layers each)
%
%  n_bi : bilateral multiplier  [1 = single, 2 = bilateral pair]
%         Head and Torso are single midline structures (n_bi = 1).
%         Arms, Hands, Legs, Feet represent BOTH sides (n_bi = 2).
%         All volumes, surface areas, TCs, and perfusion conductances
%         are multiplied by n_bi after computation for one limb.
%
%  Layer order:  1 = Core (innermost) … 4 = Skin (outermost)
%
%  —— DERIVATION OF OUTER RADII  (Table 1, finalized) ——————————————————————
%  Full derivations and primary-source citations for every layer boundary
%  are given in the manuscript Methods §2.1–2.2 (Table 1).  Summary below.
%
%  HEAD (sphere)  R_co=41.0  R_ms=46.8  R_fa=48.3  R_sk=50.05 mm
%      r_(head,core)=(r_mu^3- 〖3*V〗 _CGM/4π)^(1/3)
%      VGM=140 cm^3
%      R_eff=(abc)^(1/3)=(BPD/2*OFD/2*H/2)^(1/3)=50.05
%      BPD = 93 mm H = 95 mm OFD = 113.5 mm
%      t_skull = 1.50 mm
%      t_fat = 1.50 mm
%      t_muscle =1.50 mm
%      R_sk = R_co + 3.60 mm = 51.45 mm (skin outer boundary, consistent
%      with Vitral et al. 2018 fetal scalp imaging)
%
%  TORSO (cylinder)  R_co=43.2  R_ms=46.4  R_fa=53.4  R_sk=54.0 mm  L=190 mm
%    R_sk = AC/(2π) = 340 mm/(2π) = 54.11 mm
%           AC = 340 mm, INTERGROWTH-21st 50th %ile at 39 wk
%           (Papageorghiou et al. Lancet 2014;384:869)
%    L    = 188 mm  (crown-rump − head − neck, Table 1 finalized)
%    t_skin   = 0.72 mm
%    t_fat    = 7.0 mm
%    t_muscle = 3.20 mm  EO + IO + TA, neonatal bedside US:
%                        1.02 + 1.16 + 1.02 mm (Ofri et al. J Pediatr Surg
%                        2018;53:1588)
%    R_co = R_sk − t_skin − t_fat − t_muscle = 43.2 mm
%
%  ARMS (each arm: humerus + forearm, hand excluded)
%    R_co=4.6  R_ms=10.0  R_fa=14.50  R_sk=15.90 mm   L=125 mm
%    R_sk = MUAC/(2π) ≈ 15.9 mm (MUAC ≈ 100 mm, n=1000 term neonates,
%           Aden Yemen study PMC7568082)
%    R_co = humerus shaft radius (cortical+marrow bone core, 60/40 split,
%           Hasgall et al. 2022)
%    L    = 116 mm (humeral + forearm bone length, term fetus, hand excluded)
%
%  HANDS (each hand)
%    R_co=5.3  R_ms=6.5  R_fa=7.5  R_sk=9.50 mm   L=63 mm
%    L    = 63 mm (Malas et al. neonatal hand morphometry; cross-check
%           Merlob et al. Birth Defects 1984;20:1)
%    R_sk = AC/(2π) = 100 mm/(2π) = 16 mm
%    R_co = R_sk − t_skin − t_fat − t_muscle = 5.3 mm
%
%  LEGS (each leg: thigh + lower leg, foot excluded)
```

```
%    R_co=3.5  R_ms=17.70  R_fa=19.50  R_sk=21.00 mm   L=180 mm
%    R_sk = (thigh + calf circumference)/(2·2π) ≈ 21.0 mm
%           (Neggers et al.1995; Aden Yemen PMC7568082)
%    L    = 180 mm (femur + tibia/fibula bone length, term fetus, foot
%           excluded; Merlob et al. 1984)
%    R_co = minimum diaphysis diameter/2= 7/2 = 3.5 mm
%    t_muscle = R_sk − R_co − t_fa − t_sk = 14.2 mm
%
%  FEET (each foot)
%    R_co=3.00  R_ms=7.89  R_fa=9.89  R_sk=11.40 mm   L=78.2 mm
%    L    = 78.2 mm (neonatal foot length at term, ~7.1–8.0 cm range;
%           Nair Hospital Mumbai n=200, Bengaluru study n=500)
%    R_sk = geometric-mean cross-sectional radius from foot width/depth
%    R_co = talus bone diameter
%
%  All bone-core radii (Arms/Hands/Legs/Feet R_co) and the limb-core
%  metabolic rate qdot_co_limb = 359 W/m³ use the same cortical (60%,
%  k=295 W/m³) + marrow (40%, k=455 W/m³) composite, Hasgall et al. 2022
%  IT'IS v4.1.  See manuscript Methods Table 1 for the complete layer-
%  thickness derivation and citation list for each limb segment.
%  ──────────────────────────────────────────────────────────────────────────

Nseg = 6;
Nlay = 4;

seg_names = {'Head','Torso','Arms','Hands','Legs','Feet'};

% seg_type: 1 = sphere (head only), 2 = cylinder (all others)
seg_type = [1; 2; 2; 2; 2; 2];

% n_bi: bilateral multiplier
n_bi = [1; 1; 2; 2; 2; 2];

% Axial lengths [m]  (head: sphere — L not used in volumes/TCs)
Lseg = [0.0000;   % Head   sphere
        0.1900;   % Torso
        0.1249;   % Arms   (per single arm; bilateral handled by n_bi)
        0.0630;    % Hands
        0.1800;   % Legs
        0.0782];  % Feet

% Outer radii [m]  R_outer(segment, layer)  — describes ONE limb
R_outer = [
    0.04100, 0.04675, 0.04825, 0.05005;   % Head   (sphere)
    0.04319, 0.04639, 0.05339, 0.05411;   % Torso
    0.00463, 0.01000, 0.01450, 0.01600;   % Arms
    0.00530, 0.00650, 0.00750, 0.00950;   % Hands
    0.00350, 0.01774, 0.01953, 0.02103;   % Legs
    0.00300, 0.00789, 0.00989, 0.01139;   % Feet
];

R_inner        = zeros(Nseg, Nlay);
R_inner(:,2)   = R_outer(:,1);
R_inner(:,3)   = R_outer(:,2);
R_inner(:,4)   = R_outer(:,3);
R_seg          = R_outer(:,4);


%% =========================================================================
%  LAYER VOLUMES   V_lay  [m³]   and   SEGMENT VOLUMES   V_seg  [m³]
%
%  Single-limb values computed first; then multiplied by n_bi.
%  Head sphere:   V_shell = (4/3)π(R_out³ − R_in³)
%  Cylinders:     V       = π(R_out² − R_in²) × L
%% =========================================================================

V_lay_1 = zeros(Nseg, Nlay);   % per single limb
for s = 1:Nseg
    for l = 1:Nlay
```

```
        if seg_type(s) == 1
            V_lay_1(s,l) = (4/3)*pi*(R_outer(s,l)^3 - R_inner(s,l)^3);
        else
            V_lay_1(s,l) = pi*(R_outer(s,l)^2 - R_inner(s,l)^2)*Lseg(s);
        end
    end
end

V_seg_1 = zeros(Nseg,1);
for s = 1:Nseg
    if seg_type(s) == 1
        V_seg_1(s) = (4/3)*pi*R_seg(s)^3;
    else
        V_seg_1(s) = pi*R_seg(s)^2*Lseg(s);
    end
end

% Apply bilateral multiplier
V_lay = V_lay_1 .* n_bi;    % Nseg×Nlay  (n_bi broadcasts along columns)
V_seg = V_seg_1 .* n_bi;

%% =========================================================================
%  OUTER SURFACE AREAS   A_surf  [m²]   (per segment, bilateral included)
%
%  Head sphere:   4π R²
%  Cylinders:     2π R L   (lateral surface only; no end caps in Pennes model)
%% =========================================================================

A_surf_1 = zeros(Nseg,1);
for s = 1:Nseg
    if seg_type(s) == 1
        A_surf_1(s) = 4*pi*R_seg(s)^2;
    else
        A_surf_1(s) = 2*pi*R_seg(s)*Lseg(s);
    end
end
A_surf = A_surf_1 .* n_bi;

%% =========================================================================
%  THERMAL CONDUCTIVITIES   k_lay(segment, layer)   [W/m·K]
%  Source: IT'IS Foundation (Hasgall et al. 2022) — tissue properties database
%          v4.1, the current reference standard for computational
%          biomedical models.
%
%  Head core (k=0.565): brain+skull lumped, volume-weighted composite of
%    grey/white matter (k≈0.49–0.57) and cortical bone (k≈0.32–0.40).
%  Head muscle/fat/skin (0.495/0.210/0.370): scalp sublayers — same generic
%    muscle/fat/skin conductivities used elsewhere (see below), since the
%    skull is now part of the lumped core.
%  Torso core (k=0.510): visceral organ composite (liver, heart, kidney,
%    lung — IT'IS values cluster 0.45–0.56).
%  Limb cores, Arms/Hands/Legs/Feet (k=0.400): cortical+marrow bone composite
%    (cortical bone k≈0.32–0.40, marrow k≈0.21–0.34, IT'IS 2022).
%  Muscle (non-Head, all limbs/torso): k = 0.490 (skeletal muscle, IT'IS).
%  Fat (all segments): k = 0.210.  Skin (all segments): k = 0.370.
%% =========================================================================

k_lay = [
    0.565, 0.565, 0.320, 0.370;  % Head
    0.510, 0.490, 0.210, 0.370;  % Torso
    0.400, 0.490, 0.210, 0.370;  % Arms
    0.400, 0.490, 0.210, 0.370;  % Hands
    0.400, 0.490, 0.210, 0.370;  % Legs
    0.400, 0.490, 0.210, 0.370;  % Feet
];

%% =========================================================================
%  INTERFACE THERMAL CONDUCTANCES   TC  [W/K]
%
```

```matlab
%  k_eff at each material interface = harmonic mean of adjacent layers:
%    k_eff = 2 k₁ k₂ / (k₁ + k₂)    [series resistance at boundary]
%
%  Cylinder hollow shell (all non-head inter-layer interfaces):
%    TC = 2π k_eff L / ln(R_outer / R_inner)
%
%  Sphere solid core (head core, inner radius = 0):
%    Standard result for radial conduction in a solid sphere:
%    Q = 4π k (T_centre - T_surface) / (1/R_inner - 1/R_outer)
%    → at R_inner = 0:  TC_cm = 4π k_brain R_co
%
%  Sphere hollow shell (head muscle, fat, skin):
%    TC = 4π k_eff R_in R_out / (R_out - R_in)
%
%  Bilateral multiplier n_bi applied at end (two parallel thermal paths).
%% ========================================================================

TC_cm_1 = zeros(Nseg,1);
TC_mf_1 = zeros(Nseg,1);
TC_fs_1 = zeros(Nseg,1);

for s = 1:Nseg
    k_cm = 2*k_lay(s,1)*k_lay(s,2) / (k_lay(s,1)+k_lay(s,2));
    k_mf = 2*k_lay(s,2)*k_lay(s,3) / (k_lay(s,2)+k_lay(s,3));
    k_fs = 2*k_lay(s,3)*k_lay(s,4) / (k_lay(s,3)+k_lay(s,4));

    if seg_type(s) == 1  % —— SPHERE (head) —————————————————————————————
        % Core is a solid sphere → use solid-sphere conductance
        TC_cm_1(s) = 4*pi * k_cm * R_outer(s,1);
        % Hollow shells
        TC_mf_1(s) = 4*pi * k_mf * R_outer(s,2)*R_outer(s,3) ...
                     / (R_outer(s,3) - R_outer(s,2));
        TC_fs_1(s) = 4*pi * k_fs * R_outer(s,3)*R_outer(s,4) ...
                     / (R_outer(s,4) - R_outer(s,3));
    else                 % —— CYLINDER ———————————————————————————————————
        TC_cm_1(s) = 2*pi*k_cm*Lseg(s) / log(R_outer(s,2)/R_outer(s,1));
        TC_mf_1(s) = 2*pi*k_mf*Lseg(s) / log(R_outer(s,3)/R_outer(s,2));
        TC_fs_1(s) = 2*pi*k_fs*Lseg(s) / log(R_outer(s,4)/R_outer(s,3));
    end
end

TC_cm = TC_cm_1 .* n_bi;
TC_mf = TC_mf_1 .* n_bi;
TC_fs = TC_fs_1 .* n_bi;

%% ========================================================================
%  SKIN → AMNIOTIC FLUID CONVECTION    h_conv   [W/m²·K]
%  Values found using forced convection of a cylinder and sphere with 1
%  cm/s water speed
%% ========================================================================

h_conv = [306.8; 222.8; 426.7; 566.9; 369; 511.4];

%% ========================================================================
%  BLOOD PERFUSION   omega   [1/s]
%
%  —— DERIVATION ——————————————————————————————————————————————————————————
%  Segment blood flows are set so that body segments + placenta = CVO:
%
%    CVO = 1544 mL/min  (Zhang and Lindsey)
%
%  CVO PARTITION  (% of CVO per bilateral group):
%    Head                   22%  (20–25% range)  — priority cerebral circulation
%    Torso                  34%  (30–35% range)  — visceral organs, lungs, coronary
%    Each arm+hand pair      3%  (3–7%  bilateral upper limbs = 6% total)
%    Each leg+foot pair      9%  (15–20% bilateral lower limbs = 18% total)
%    Placenta               20%                  — umbilical vein (Gill 1981)
%    Total                 100%  ✓
%
```

```
%  Arms and hands are separate model segments but form one upper limb unit
%  (3% per side).  Their individual flows are split by segment volume:
%    Q_arm  = 3% × V_arm  / (V_arm+V_hand)  per side  → bilateral = 6% × frac_arm
%    Q_hand = 3% × V_hand / (V_arm+V_hand)  per side  → bilateral = 6% × frac_hand
%  Leg and foot follow the same logic (9% per side, split by volume):
%    frac_arm  = V_arm  / (V_arm+V_hand)  ≈ 79%
%    frac_hand = V_hand / (V_arm+V_hand)  ≈ 21%
%    frac_leg  = V_leg  / (V_leg+V_foot)  ≈ 83%
%    frac_foot = V_foot / (V_leg+V_foot)  ≈ 17%
%
%  WITHIN-SEGMENT LAYER FRACTIONS (fraction of segment total flow):
%    Head:  core 80% (WM+subcortex), muscle 12% (cortical GM),
%           fat 4% (skull), skin 4% (scalp)
%           → omega_co ~86 mL/min/100g  (cerebral blood flow; literature ~80–120)
%    Torso: core 90% (liver, kidney, heart, lung dominate),
%           muscle 4%, fat 3%, skin 3%
%           → omega_co ~30 mL/min/100g  (mixed visceral; literature ~30–60)
%    Limbs: core 5% (bone marrow), muscle 85% (dominant limb perfusion),
%           fat 5%, skin 5%
%           → omega_mu ~40–100 mL/min/100g
%
%  References:
%    Rudolph AM. Circ Res 1985;57(6):811–821.  (CVO partition; carcass flows)
%    Hasgall PA et al. IT'IS v4.1 2022.         (tissue densities)
%  ──────────────────────────────────────────────────────────────────────────

% —— CVO and target bilateral flows  [mL/min] ─────────────────────────────
BW_kg      = 3.2;
CVO_ml     = 1544;                         % 1544 mL/min (Zhang and Lindsey)
Q_plac_ml  = 0.2*CVO_ml;

% Volume-proportional split of limb flow into individual segments
frac_arm   = V_seg_1(3) / (V_seg_1(3) + V_seg_1(4));   % ~0.79
frac_hand  = V_seg_1(4) / (V_seg_1(3) + V_seg_1(4));   % ~0.21
frac_leg   = V_seg_1(5) / (V_seg_1(5) + V_seg_1(6));   % ~0.83
frac_foot  = V_seg_1(6) / (V_seg_1(5) + V_seg_1(6));   % ~0.17

% % CVO per bilateral segment  (Mielke & Benda 2001: 21.7/33.7/5.7/17.7)
pct_CVO = [21.7;                  % Head
           33.7;                  % Torso
           5.7 * frac_arm;        % Arms   (2.85% per arm+hand side, volume-split)
           5.7 * frac_hand;       % Hands
           17.7 * frac_leg;       % Legs   (8.85% per leg+foot side, volume-split)
           17.7 * frac_foot];     % Feet

Q_seg_target = pct_CVO / 100 * CVO_ml;   % mL/min bilateral

% —— Layer fractions [core, muscle, fat, skin] per segment ────────────────
layer_frac = [
    0.80, 0.12, 0.04, 0.04;    % Head   (WM/subcortex, GM, skull, scalp)
    0.90, 0.04, 0.03, 0.03;    % Torso  (viscera dominate)
    0.05, 0.85, 0.05, 0.05;    % Arms   (bone, muscle, fat, skin)
    0.05, 0.80, 0.08, 0.07;    % Hands  (metacarpal, intrinsic mu, fat, skin)
    0.05, 0.85, 0.05, 0.05;    % Legs   (femur/tibia, thigh/calf, fat, skin)
    0.05, 0.75, 0.10, 0.10;    % Feet   (tarsal, plantar, fat, skin)
];

% —— Compute layer omega = Q_layer / (V_layer × 60)  [1/s] ───────────────
%  Q_layer [mL/min] = layer_frac × Q_seg_target  (bilateral)
%  V_layer [mL]     = V_lay(i,l) × 1e6           (bilateral volume)
omega_co = zeros(Nseg,1);
omega_mu = zeros(Nseg,1);
omega_fa = zeros(Nseg,1);
omega_sk = zeros(Nseg,1);

for i = 1:Nseg
    omega_co(i) = (layer_frac(i,1) * Q_seg_target(i) / 60) / (V_lay(i,1) * 1e6);
    omega_mu(i) = (layer_frac(i,2) * Q_seg_target(i) / 60) / (V_lay(i,2) * 1e6);
    omega_fa(i) = (layer_frac(i,3) * Q_seg_target(i) / 60) / (V_lay(i,3) * 1e6);
```

```
    omega_sk(i) = (layer_frac(i,4) * Q_seg_target(i) / 60) / (V_lay(i,4) * 1e6);
end

% Volume-weighted segment omega (for reference; not used directly in solver)
omega = (omega_co.*V_lay_1(:,1) + omega_mu.*V_lay_1(:,2) + ...
         omega_fa.*V_lay_1(:,3) + omega_sk.*V_lay_1(:,4)) ./ V_seg_1;

% ── Perfusion conductance  Φ = ω ρ_b c_b V  [W/K] ───────────────────────
Phi = zeros(Nseg, Nlay);
Phi(:,1) = omega_co .* rho_b .* c_b .* V_lay(:,1);
Phi(:,2) = omega_mu .* rho_b .* c_b .* V_lay(:,2);
Phi(:,3) = omega_fa .* rho_b .* c_b .* V_lay(:,3);
Phi(:,4) = omega_sk .* rho_b .* c_b .* V_lay(:,4);

%% =========================================================================
%  METABOLIC HEAT GENERATION   qdot  [W/m³]
%
%  ── DERIVATION ────────────────────────────────────────────────────────────
%  Formula:  qdot = Ki × ρ
%  where Ki = specific metabolic rate [W/kg] and ρ = tissue density [kg/m³].
%
%  Ki source: Elia M. "Organ and tissue contribution to metabolic rate."
%             In: Kinney JM, Tucker HN (eds). Energy metabolism: tissue
%             determinants and cellular corollaries.
%             Raven Press, New York 1992:61–80 (CIBA Symp 164).
%  Validated: Wang ZM et al. Am J Clin Nutr 2010;92:1369 (n=131, MRI+calorimetry)
%
%  Density source: IT'IS Foundation (Hasgall et al. 2022)
%
%  Ki conversion:  1 kcal/kg/day × 4184 J/kcal ÷ 86400 s/day = 0.04843 W/kg
%
%  UNIFORM OUTER LAYERS (same for all segments):
%    Muscle: Ki = 13 kcal/kg/day × 1090 kg/m³ × 0.04843 = 686 W/m³
%    Fat:    Ki = 4.5 × 911 × 0.04843               = 199 W/m³  ← 3.4× Fiala
%    Skin:   Ki = 12 × 1109 × 0.04843               = 644 W/m³  ← 1.75× Fiala
%    (Fiala used 58 and 368 W/m³ respectively; Elia/IT'IS values are correct)
%
%  SEGMENT-SPECIFIC CORE qdot:
%    Head core (brain + skull, LUMPED — Table 1 finalized geometry,
%    R_co = 47.85 mm, V_co = 458.9 cm³ = brain 364.9 cm³ + skull 94.0 cm³):
%      Q_brain = 240 kcal/kg/day × 0.04843 W/kg/(kcal/kg/day) × 0.362 kg
%              = 4.43 W
%        (Elia 1992 whole-brain Ki; brain mass 0.362 kg from Andescavage 2017)
%      Skull bone contributes negligible metabolic heat at this Ki and is
%      treated as part of the inert core volume only (qdot from brain alone).
%        qdot_co(head) = Q_brain / V_co = 4.43 / 288.7e-6 ≈ 15,553 W/m³
%      Outer 3 layers = scalp sublayers (muscle/fat/skin, 1.20 mm each;
%      Erim et al. 2024 FST=3.6mm; Vitral 2018 skin boundary), using the
%      generic muscle/fat/skin qdot below (uniform outer layers).
%
%    Torso core (visceral organ mix, R_co = 44.11 mm, L = 188 mm,
%    V_co = π R_co² L ≈ 1048 cm³):
%      Q_organs = Σ Ki_organ × m_organ
%      Organ masses (Gruenwald P. Am J Obstet Gynecol 1966;94:1112; BW=3200g):
%        liver 141.4g, heart 21.34g, kidney 26.78g, lung 57.8g, spleen 10.82g
%      Elia Ki: liver=200, heart=440, kidney=440, lung=12, spleen=12 kcal/kg/d
%      Q_organs=(200*0.1414+440*0.02134+440*0.02678+12*0.0578+12*0.01082)*0.04843
%              =2.435 W
%        qdot_co(torso) = Q_organs / V_co = 2.435 / 1048e-6 ≈ 2,322 W/m³
%
%    Limb cores (Arms, Hands, Legs, Feet — bone: 60% cortical + 40% marrow,
%    Hasgall et al. 2022 IT'IS v4.1):
%        qdot_cortical = 3 kcal/kg/d × 2031 kg/m³ × 0.04843 = 295 W/m³
%        qdot_marrow   = 12 kcal/kg/d × 1027 kg/m³ × 0.04843 = 597 W/m³ →
%                        rounded to 455 W/m³ (yellow/red marrow composite,
%                        IT'IS v4.1 marrow Ki)
%        qdot_co_limb  = 0.60×295 + 0.40×455 = 359 W/m³
%
%  TOTAL MODEL BMR (at 39 wk, BW=3.2 kg, finalized Table 1 geometry):
```

```
%      Target: 7.6 W from Fick principle
%              (Acharya & Sitras Acta Obstet Gynecol Scand 2009;88:104;
%               Fumia et al. Am J Obstet Gynecol 1984;150:549)
%      Model:  7.84 W (+3.2%) — Head 4.49 W, Torso 2.91 W, limbs 0.45 W
% ──────────────────────────────────────────────────────────────────────────

qdot_co = [15553; 2322;  394;  394;  394;  394];        % Head core: brain 4.43W/V_co=288.7cm3
(Elia 1992 + Andescavage 2017); Torso: 2.435W/1048.43cm3 (Gruenwald 1966); limbs: 0.6x295+0.4x455
(Hasgall et al. 2022)
qdot_mu = [  686;  686;  686;  686;  686;  686];        % Uniform skeletal muscle (Elia 1992);
head = face/temporal muscle
qdot_fa = [  199;  199;  199;  199;  199;  199];        % Adipose (Elia 1992 / IT'IS)
qdot_sk = [  644;  644;  644;  644;  644;  644];        % Skin    (Elia 1992 / IT'IS)

%% =========================================================================
%  UTERINE WALL BOUNDARY CONDITION
%
%  —— DERIVATION ───────────────────────────────────────────────────────────
%  The fetal skin is separated from the maternal uterine wall by the
%  amniotic fluid.  Heat flows: T_af → amniotic fluid → myometrium → T_m.
%
%  Two resistances in series:
%    1. Amniotic fluid convection:  h_af = 17–50 W/m²K (natural convection)
%    2. Myometrium wall conduction: t_myo ≈ 10 mm, k_myo ≈ 0.50 W/mK
%       → R_myo = t/k = 0.01/0.50 = 0.020 m²K/W → h_myo = 50 W/m²K
%    Combined: 1/h_total = 1/17 + 1/50 → h_total ≈ 12.8 ≈ 13 W/m²K
%    The myometrium accounts for ~83% of total resistance.
%
%
%  CONTRAST WITH v1: h=50, A=0.50 → h×A = 25 W/K (5× too large),
%  which over-cooled T_af to only +0.09°C above T_m.
% ──────────────────────────────────────────────────────────────────────────

h_uterus = 22;  % combined uterine conductance     [W/m²·K]
A_uterus = 0.15;   % inner uterine wall area           [m²]

%% =========================================================================
%  UMBILICAL CORD + PLACENTA  (NTU-effectiveness, counterflow model)
%
%  —— DERIVATION ───────────────────────────────────────────────────────────
%  BLOOD FLOW:
%    Measured by pulsed Doppler ultrasound at term:
%    Q_UV = 318 mL/min (umbilical vein flow per kg fetal weight)
%    Source: Zhang and Lindsey
%    Confirmed: Konje JC et al. Br J Obstet Gynaecol 1996 (transit-time
%               flowmeter at C/S: ~90 mL/min/kg); Acharya & Sitras
%               Acta Obstet Gynecol Scand 2016;95:1107 (review: 64–90 mL/kg)
%
%     mUA = ρ_blood × Q_UV
%        = 1060 kg/m³ × (318×10⁻⁶ m³/min / 60 s/min)
%        = 1060 × 5.30×10⁻⁶ m³/s
%        = 5.62×10⁻³ kg/s
%
%  PLACENTAL HEAT EXCHANGE EFFECTIVENESS:
%    ε = 0.845 (84.5% of fetal heat exits via placenta)
%    Source: Gilbert RD et al. J Appl Physiol 1985;59:634 — ten
%    ewe/fetal-lamb dyads, direct calorimetry.
%    Consistent with clinical observation (Schröder & Power 1997 review):
%    80–85% placenta, 15–20% via amniotic fluid conduction.
%
% ──────────────────────────────────────────────────────────────────────────

Q_UV_m3s= (318*10^-6)/60; % m^3/s 318 mL/min
mUA =rho_b*Q_UV_m3s;         % 0.00556 umbilical arterial mass flow (kg/s)

%  —— UMBILICAL CORD + PLACENTA (Ec/Ep effectiveness formulation) ──────────
%
%  Ec = 0.32   cord counterflow heat exchanger effectiveness
%               (fraction of maximum possible cord heat exchange)
```

```
%  Ep = 0.845  placenta heat pathway partition
%
%  Z5 = fraction of arterial blood temperature returned to fetal pool
%  Z6 = fraction coupling T_a to T_m (drives placental heat removal)
%
%  umb = mUA × c_b × Z6   [W/K]  effective umbilical conductance

Ec = 0.32;    % cord counterflow effectiveness
Ep = 0.845;   % placenta heat pathway partition fraction (Gilbert 1985)

Z5      = Ec + ((1-Ep)*(1-Ec)^2) / (1 - Ec*(1-Ep));
Z6_coef = (Ep*(1-Ec))            / (1 - Ec*(1-Ep));
umb     = mUA * c_b * Z6_coef;   % effective umbilical coupling  [W/K]

%% =========================================================================
%  SINGLE-POINT SOLVE  (Tmother = 37.0 °C)
%% =========================================================================

res = solve_26x26(Tmother, Nseg, Nlay, Phi, TC_cm, TC_mf, TC_fs, ...
                  h_conv, A_surf, qdot_co, qdot_mu, qdot_fa, qdot_sk, ...
                  V_lay, h_uterus, A_uterus, umb);

Ta  = res.Ta;
Taf = res.Taf;

fprintf('\n%s\n', repmat('=',1,72));
fprintf('  BABY STOLWIJK 26×26 v3  —  Single-point results\n');
fprintf('%s\n\n', repmat('=',1,72));
fprintf('  Tmother = %.2f °C\n', Tmother);
fprintf('  omega_UV    = %.1f mL/min  (%.0f mL/min/kg)\n', ...
        Q_UV_m3s*1e6*60, Q_UV_m3s*1e6*60/3.2);
fprintf('  mUA     = %.5f kg/s,   C_min = %.2f W/K\n', mUA, mUA*c_b);
fprintf('  Ec = %.3f (cord)   Ep = %.3f (placenta partition)\n', Ec, Ep);
fprintf('  Ta  (central blood) = %.4f °C  (+%.4f)\n', Ta,  Ta-Tmother);
fprintf('  Taf (amniotic)      = %.4f °C  (+%.4f)\n', Taf, Taf-Tmother);
fprintf('  h×A (uterine)       = %.3f W/K\n\n', h_uterus*A_uterus);

fprintf('  %-8s %8s %10s %8s %8s\n', 'Segment','T_core','T_muscle','T_fat','T_skin');
fprintf('  %s\n', repmat('-',1,50));
for i = 1:Nseg
    fprintf('  %-8s %8.4f %10.4f %8.4f %8.4f\n', seg_names{i}, ...
            res.T_co(i), res.T_mu(i), res.T_fa(i), res.T_sk(i));
end

Q_total = sum(qdot_co.*V_lay(:,1) + qdot_mu.*V_lay(:,2) + ...
              qdot_fa.*V_lay(:,3) + qdot_sk.*V_lay(:,4));
fprintf('\n  Total model BMR = %.3f W  \n\n', Q_total);

%% —— BLOOD PERFUSION PER SEGMENT ——————————————————————————————————————————
%  All layer omega values are computed directly from CVO % targets and
%  physiological layer fractions (see BLOOD PERFUSION section above).
%  omega_co is NO LONGER back-calculated.
%% —————————————————————————————————————————————————————————————————————————

% Bilateral layer flows [mL/min]
Q_seg_flow = Q_seg_target;                       % from pct_CVO above
Q_co_flow  = omega_co .* V_lay(:,1) * 1e6 * 60;
Q_mu_flow  = omega_mu .* V_lay(:,2) * 1e6 * 60;
Q_fa_flow  = omega_fa .* V_lay(:,3) * 1e6 * 60;
Q_sk_flow  = omega_sk .* V_lay(:,4) * 1e6 * 60;

fprintf('  BLOOD PERFUSION — segment level  [mL/min]\n');
fprintf('  %s\n', repmat('-',1,68));
fprintf('  %-8s  %10s  %10s  %10s  %10s  %10s\n', ...
        'Segment','Q_total','Q_core','Q_muscle','Q_fat','Q_skin');
fprintf('  %s\n', repmat('-',1,68));
for i = 1:Nseg
    fprintf('  %-8s  %10.2f  %10.2f  %10.2f  %10.2f  %10.2f\n', ...
            seg_names{i}, Q_seg_flow(i), Q_co_flow(i), ...
```

```matlab
            Q_mu_flow(i), Q_fa_flow(i), Q_sk_flow(i));
end
fprintf('  %s\n', repmat('-',1,68));
fprintf('  %-8s  %10.2f  %10.2f  %10.2f  %10.2f  %10.2f\n', ...
        'TOTAL', sum(Q_seg_flow), sum(Q_co_flow), ...
        sum(Q_mu_flow), sum(Q_fa_flow), sum(Q_sk_flow));
fprintf('  %s\n\n', repmat('-',1,68));

% CVO balance check — body segments + placenta = CVO exactly
Q_CVO_est   = 450 * 3.2;          % mL/min  (450 mL/min/kg × BW, Rudolph 1985)
Q_uv        = Q_UV_m3s * 1e6 * 60;% mL/min  umbilical vein (placenta)
fprintf('  CVO BALANCE  [mL/min]\n');
fprintf('  %s\n', repmat('-',1,48));
for i = 1:Nseg
    fprintf('  %-8s                    %8.2f\n', seg_names{i}, Q_seg_flow(i));
end
fprintf('  %s\n', repmat('-',1,48));
fprintf('  %-28s  %8.2f\n', 'Body segments subtotal',   sum(Q_seg_flow));
fprintf('  %-28s  %8.2f\n', 'Placenta (umbilical vein)', Q_uv);
fprintf('  %s\n', repmat('-',1,48));
fprintf('  %-28s  %8.2f\n', 'TOTAL  (= CVO)',  sum(Q_seg_flow) + Q_uv);


%% =========================================================================
%  TEMPERATURE SWEEPS
%% =========================================================================

% Fig A: 36.5 – 41.0 °C (Guenkawa Fig 7 equivalent)
Tm_A  = (365:410) * 0.1;
nA    = numel(Tm_A);
resA  = zeros(nA, 4);    % [T_co_head, T_sk_head, Ta, Taf]
for i = 1:nA
    ri = solve_26x26(Tm_A(i), Nseg, Nlay, Phi, TC_cm, TC_mf, TC_fs, ...
                     h_conv, A_surf, qdot_co, qdot_mu, qdot_fa, qdot_sk, ...
                     V_lay, h_uterus, A_uterus, umb);
    resA(i,:) = [ri.T_co(1), ri.T_sk(1), ri.Ta, ri.Taf];
end

% Fig B: Lavesson comparison
%   Lavesson (2017) measured maternal AXILLARY temperature (not core).
%   Axillary ≈ T_m − 0.27°C (Lefrant 2003 Intensive Care Med 29:414).
%   Model x-axis converted to axillary by subtracting 0.27.
%   Model y-axis = T_sk(head) (intradermal scalp ~2mm ≈ skin node).

dT_ax    = 0.27;
Tm_B_co  = (349:399)*0.1 + dT_ax;    % maternal CORE passed to solver
Tm_B_ax  = Tm_B_co - dT_ax;          % maternal AXILLARY for x-axis
nB       = numel(Tm_B_co);
Tsk_B    = zeros(nB,1);
Tsk_GUENKAWA = zeros(nB,1);
for i = 1:nB
    ri      = solve_26x26(Tm_B_co(i), Nseg, Nlay, Phi, TC_cm, TC_mf, TC_fs, ...
                          h_conv, A_surf, qdot_co, qdot_mu, qdot_fa, qdot_sk, ...
                          V_lay, h_uterus, A_uterus, umb);
    Tsk_B(i) = ri.T_sk(1);
    Tsk_GUENKAWA2=UniformModelFunction(Tm_B_co(i));
    Tsk_GUENKAWA(i) = Tsk_GUENKAWA2(4);
end

% Load Lavesson scatter
try
    lav = readmatrix('lavesson_full_data.xlsx');
    Tm_lav = lav(:,1);  Tf_lav = lav(:,2);
    have_lav = true;
catch
    warning('lavesson_full_data.xlsx not found — scatter omitted from Fig B.');
    have_lav = false;
end
```

```matlab
% Fig C: radial profiles at three T_m values
Tma   = [36.5, 37.2, 39.0];
resC  = cell(1,3);
for i = 1:3
    resC{i} = solve_26x26(Tma(i), Nseg, Nlay, Phi, TC_cm, TC_mf, TC_fs, ...
                          h_conv, A_surf, qdot_co, qdot_mu, qdot_fa, qdot_sk, ...
                          V_lay, h_uterus, A_uterus, umb);
end


%% ========================================================================
%  GLOBAL PLOT DEFAULTS — set BEFORE any figure is created
%
%  MATLAB dark theme injects groot-level defaults that override per-figure
%  Color properties (axes background goes dark, text goes light).
%  The only reliable fix is to reset every relevant groot default here so
%  the theme has nothing left to override.
%% ========================================================================

% Typography
set(groot, 'defaultAxesFontName',       'Arial');
set(groot, 'defaultTextFontName',       'Arial');
set(groot, 'defaultAxesFontSize',       9);
set(groot, 'defaultTextFontSize',       9);

% Line widths
set(groot, 'defaultLineLineWidth',      0.75);
set(groot, 'defaultAxesLineWidth',      0.75);

% —— Colour overrides that defeat dark theme ——————————————————————————————
set(groot, 'defaultFigureColor',        [1 1 1]);   % figure window: white
set(groot, 'defaultAxesColor',          [1 1 1]);   % axes plot area: white
set(groot, 'defaultAxesXColor',         [0 0 0]);   % x axis, ticks, tick labels
set(groot, 'defaultAxesYColor',         [0 0 0]);   % y axis, ticks, tick labels
set(groot, 'defaultAxesZColor',         [0 0 0]);
set(groot, 'defaultTextColor',          [0 0 0]);   % all text objects: black
set(groot, 'defaultAxesGridColor',      [0.85 0.85 0.85]);
set(groot, 'defaultAxesMinorGridColor', [0.92 0.92 0.92]);

%% ========================================================================
%  FIGURE A — Head skin temperature vs maternal axillary + Lavesson
%
%  CORRECTION vs v1:
%    v1 plotted T_co(head) vs T_m (maternal CORE) — double mismatch of ~0.3°C
%    v3 plots  T_sk(head) vs T_m−0.27°C (maternal AXILLARY) — correct pairing
%% ========================================================================

hA = figure('Name','FigA_LavessonComparison', 'Color',[1 1 1], 'Visible','on');
hA_num = hA.Number;
apply_pubstyle(hA, 7.5, 4.8);
set(groot, 'defaultAxesTickDir', 'in');
set(gca,'TickDir','in','Box','on');
cMulti   = [0 176 240]/255;
cUniform  = [255 192 0]/255;

plot(Tm_B_ax, Tsk_B,'Color',cMulti, 'LineStyle', '-', 'LineWidth', 1.2, ...
     'DisplayName','multilayer model');  hold on;
plot(Tm_B_ax, Tsk_GUENKAWA,'Color',cUniform, 'LineStyle','--', 'LineWidth', 1.2, ...
     'DisplayName','6-cylinder model');  hold on;
gray=[0.5, 0.5, 0.5];
if have_lav
    lav=plot(Tm_lav, Tf_lav, 'o', ...
         'MarkerEdgeColor',gray,'MarkerFaceColor',gray, ...
         'MarkerSize',3,'LineWidth',0.75, ...
         'DisplayName','Lavesson measurements');
end
fontname("Arial");
xlabel('Maternal axillary temperature (°C)','FontSize', 12);
ylabel('Fetal scalp temperature (°C)','FontSize', 12);
```

```
xlim([35 40]);  ylim([35 40]);
legend(lav,'Lavesson measurements','Location','SouthWest','Box','off','FontSize', 12);
apply_pubstyle(hA, 7.5, 4.8);
set(gca,'TickDir','in','Box','on');
drawnow;
drawnow;
figure(hA_num);
print('-djpeg', '-r600', fullfile(out_dir,'FigA_LavessonComparison.jpg'));

%% ==========================================================================
%  FIGURE B — Per-segment temperature profiles  (2 × 3 panel)
%
%  Each subplot shows one segment at Tmother = 37.0 °C.
%  X-axis layout (normalised):
%    [0 … 1]   = within-segment radial profile  (core → muscle → fat → skin)
%    gap at 1.0 – 1.15
%    x = 1.15  = T_af  (amniotic fluid – just outside the skin surface)
%    x = 1.30  = T_m   (maternal core  – ultimate heat sink)
%  This makes the full thermal drop from fetal core to maternal surroundings
%  visible in a single chart per segment.
%
%  Markers:
%    ● filled circle   – layer nodal temperatures  (T_co, T_mu, T_fa, T_sk)
%    ◆ filled diamond  – T_af
%    ▼ filled triangle – T_m  (reference)
%    Dashed grey line bridges T_sk → T_af → T_m.
%    Dotted red line marks the T_m level across each subplot.
%    Dotted vertical grey line marks the skin boundary at r/R = 1.
%% ==========================================================================

r37  = res;          % single-point result at 37 °C (solved above)
Taf37 = r37.Taf;

x_af = 1.15;         % x position for T_af marker
x_tm = 1.30;         % x position for T_m  marker

% Mid-radius (normalised) for each layer
R_mid_B = zeros(Nseg, Nlay);
for s = 1:Nseg
    R_mid_B(s,1) = R_outer(s,1) / 2;
    for l = 2:Nlay
        R_mid_B(s,l) = (R_inner(s,l) + R_outer(s,l)) / 2;
    end
end

hB = figure('Name','FigB_PerSegmentProfiles','Color',[1 1 1],'Visible','on');
hB_num = hB.Number;
apply_pubstyle(hB, 7.5, 4.8);

for s = 1:Nseg
    axs = subplot(2, 3, s);

    T_nodes = [r37.T_co(s); r37.T_mu(s); r37.T_fa(s); r37.T_sk(s)];
    rN      = R_mid_B(s,:)' ./ R_seg(s);

    % Within-segment line (centre → skin)
    r_line = [0; rN; 1.0];
    T_line = [T_nodes(1); T_nodes; T_nodes(4)];
    plot(r_line, T_line, '-k', 'LineWidth', 1.3);  hold on;

    % Dashed connector: skin → T_af → T_m
    plot([1.0, x_af, x_tm], [T_nodes(4), Taf37, Tmother], ...
         '--', 'Color', [0.45 0.45 0.45], 'LineWidth', 0.9);

    % Node markers (filled circles)
    plot(rN, T_nodes, 'ok', ...
         'MarkerFaceColor', [0 0 0], 'MarkerSize', 5, 'LineWidth', 0.5);

    % T_af marker (filled blue diamond)
```

```
    plot(x_af, Taf37, 'd', ...
         'Color', [0.15 0.15 0.80], 'MarkerFaceColor', [0.15 0.15 0.80], ...
         'MarkerSize', 5);

    % T_m marker (filled red downward triangle)
    plot(x_tm, Tmother, 'v', ...
         'Color', [0.80 0.10 0.10], 'MarkerFaceColor', [0.80 0.10 0.10], ...
         'MarkerSize', 5);

    % T_m horizontal reference line
    yl = ylim;
    plot([0, x_tm+0.06], [Tmother Tmother], ':', ...
         'Color', [0.80 0.10 0.10], 'LineWidth', 0.7);

    % Vertical separator at r/R = 1 (skin boundary)
    yl2 = ylim;
    plot([1.0 1.0], yl2, ':', 'Color', [0.70 0.70 0.70], 'LineWidth', 0.7);

    % Temperature annotations next to each node
   xOffsets = [0.04, 0.04, 0.04, 0.04];
   yOffset = 0.01;

for l = 1:Nlay
    text(rN(l) + xOffsets(l), T_nodes(l) + yOffset, ...
         sprintf('%.3f', T_nodes(l)), ...
         'FontSize', 6.5, ...
         'Color', [0 0 0], ...
         'VerticalAlignment', 'bottom', ...
         'FontName', 'Arial');
end
    text(x_af + 0.03, Taf37, sprintf('%.3f', Taf37), ...
         'FontSize', 6.5, 'Color', [0.10 0.10 0.75], ...
         'VerticalAlignment', 'middle', 'FontName', 'Arial');
    text(x_tm + 0.03, Tmother + 0.01, sprintf('%.1f', Tmother), ...
         'FontSize', 6.5, 'Color', [0.75 0.05 0.05], ...
         'VerticalAlignment', 'bottom', 'FontName', 'Arial');

    % Axis formatting
    xlim([0, x_tm + 0.10]);
    set(axs, ...
        'XTick',      [0, 0.5, 1.0, x_af, x_tm], ...
        'XTickLabel', {'0', '0.5', '1', 'T_{af}', 'T_m'}, ...
        'Color',  [1 1 1], 'XColor', [0 0 0], 'YColor', [0 0 0], ...
        'FontName', 'Arial', 'FontSize', 8, ...
        'Box', 'on', 'TickDir', 'out', 'LineWidth', 0.8);
    axs.XLabel.Color = [0 0 0];
    axs.YLabel.Color = [0 0 0];
    axs.Title.Color  = [0 0 0];
    xlabel('r / R   →   boundary', 'FontSize', 8);
    ylabel('Temperature (°C)', 'FontSize', 8);
    title(seg_names{s}, 'FontWeight', 'normal', 'FontSize', 9);
end

% Shared legend using invisible axes at the bottom
ax_leg = axes('Position', [0.25 0.005 0.55 0.055], ...
              'Visible', 'off', 'Color', [1 1 1], ...
              'XColor', [1 1 1], 'YColor', [1 1 1]);
hold(ax_leg, 'on');
h1 = plot(ax_leg, NaN, NaN, '-ok', 'MarkerFaceColor', 'k', 'MarkerSize', 4);
h2 = plot(ax_leg, NaN, NaN, '--',  'Color', [0.45 0.45 0.45], 'LineWidth', 0.9);
h3 = plot(ax_leg, NaN, NaN, 'd',   'Color', [0.15 0.15 0.80], ...
          'MarkerFaceColor', [0.15 0.15 0.80], 'MarkerSize', 4);
h4 = plot(ax_leg, NaN, NaN, 'v',   'Color', [0.80 0.10 0.10], ...
          'MarkerFaceColor', [0.80 0.10 0.10], 'MarkerSize', 4);
h5 = plot(ax_leg, NaN, NaN, ':',   'Color', [0.80 0.10 0.10], 'LineWidth', 0.7);
lg_d = legend(ax_leg, [h1 h2 h3 h4 h5], ...
              {'Layer nodes  (T_{co} → T_{sk})', 'Connector to boundary', ...
               'T_{af}  (amniotic fluid)', 'T_m  (maternal core)', ...
               'T_m  reference line'}, ...
```

```
               'Orientation', 'horizontal', 'Box', 'off', ...
               'FontSize', 7, 'FontName', 'Arial', 'TextColor', [0 0 0]);
lg_d.Color     = [1 1 1];
lg_d.TextColor = [0 0 0];

apply_pubstyle(hB, 7.5, 4.8);   % re-apply after all children created
drawnow;
drawnow;
figure(hB_num);
print('-djpeg', '-r600', fullfile(out_dir,'FigB_PerSegmentProfiles.jpg'));
fprintf('Figure B saved.\n');

%% ========================================================================
%  EXCEL OUTPUT  —  BabyStolwijk_temperatures.xlsx
%  Sheet 1: Single_Point_37C   — all temperatures at T_m = 37 °C
%  Sheet 2: Temperature_Sweep  — full sweep over Fig B range
%  Sheet 3: Checks_and_Balances — heat routing, energy check, CVO
%% ========================================================================

layer_lbl = {'Core','Muscle','Fat','Skin'};
fname_xls = fullfile(out_dir, 'BabyStolwijk_temperatures.xlsx');

% —— Sheet 1: single point (T_m = 37 °C) ——————————————————————————————————
param_names = { ...
    'T_maternal_core_C';     ...
    'T_maternal_axillary_C'; ...   % T_m − 0.27 (Lefrant 2003)
    'T_amniotic_fluid_C';    ...
    'T_fetal_scalp_skin_C';  ...   % T_sk(Head) — closest to Lavesson sensor
    'T_central_blood_C'};

param_vals = [Tmother; Tmother-dT_ax; res.Taf; res.T_sk(1); res.Ta];

for s = 1:Nseg
    for l = 1:Nlay
        param_names{end+1} = sprintf('T_%s_%s_C', layer_lbl{l}, seg_names{s}); %#ok<SAGROW>
        param_vals(end+1)  = res.Temps(4*(s-1)+l);                              %#ok<SAGROW>
    end
end

T_sp = table(param_names, param_vals, param_vals - Tmother, ...
    'VariableNames', {'Parameter','Temperature_C','Delta_from_Tm_C'});
writetable(T_sp, fname_xls, 'Sheet', 'Single_Point_37C');

% —— Sheet 2: temperature sweep (Fig B range) ——————————————————————————————
Taf_sw  = zeros(nB,1);
Tsk_sw  = zeros(nB,1);
Ta_sw   = zeros(nB,1);
Tseg_sw = zeros(nB, Nseg*Nlay);

for i = 1:nB
    ri = solve_26x26(Tm_B_co(i), Nseg, Nlay, Phi, TC_cm, TC_mf, TC_fs, ...
        h_conv, A_surf, qdot_co, qdot_mu, qdot_fa, qdot_sk, ...
        V_lay, h_uterus, A_uterus, umb);
    Taf_sw(i) = ri.Taf;  Tsk_sw(i) = ri.T_sk(1);  Ta_sw(i) = ri.Ta;
    for s = 1:Nseg
        Tseg_sw(i,(s-1)*Nlay+1) = ri.T_co(s);
        Tseg_sw(i,(s-1)*Nlay+2) = ri.T_mu(s);
        Tseg_sw(i,(s-1)*Nlay+3) = ri.T_fa(s);
        Tseg_sw(i,(s-1)*Nlay+4) = ri.T_sk(s);
    end
end

sw_hdr = {'T_maternal_core_C','T_maternal_axillary_C', ...
    'T_amniotic_fluid_C','T_fetal_scalp_skin_C','T_central_blood_C'};
for s = 1:Nseg
    for l = 1:Nlay
        sw_hdr{end+1} = sprintf('T_%s_%s_C', layer_lbl{l}, seg_names{s}); %#ok<SAGROW>
    end
end
```

```
T_sw = array2table([Tm_B_co', Tm_B_ax', Taf_sw, Tsk_sw, Ta_sw, Tseg_sw], ...
    'VariableNames', sw_hdr);
writetable(T_sw, fname_xls, 'Sheet', 'Temperature_Sweep');

% —— Sheet 3: Checks & Balances ——————————————————————————————————————————
dT_scalp_core_sw = Tsk_sw - Tm_B_co';
dT_scalp_ax_sw   = Tsk_sw - Tm_B_ax';
dT_af_core_sw    = Taf_sw - Tm_B_co';
dT_af_ax_sw      = Taf_sw - Tm_B_ax';

Q_wall_sw  = h_uterus * A_uterus * dT_af_core_sw;
Q_plac_sw  = umb * (Ta_sw - Tm_B_co');
Q_check_sw = Q_wall_sw + Q_plac_sw;
Q_wall_pct = Q_wall_sw / Q_total * 100;
Q_plac_pct = Q_plac_sw / Q_total * 100;

Q_UV_mLmin_val  = Q_UV_m3s * 1e6 * 60;
Q_CVO_mLmin_val = 450 * BW_kg;
umb_pct_val     = Q_UV_mLmin_val / Q_CVO_mLmin_val * 100;

cb3_hdr = {'T_maternal_core_C','T_maternal_axillary_C', ...
    'T_amniotic_fluid_C','T_fetal_scalp_skin_C','T_central_blood_C', ...
    'dT_scalp_minus_Tcore_C','dT_scalp_minus_Taxillary_C', ...
    'dT_Taf_minus_Tcore_C','dT_Taf_minus_Taxillary_C', ...
    'Q_BMR_total_W','Q_skin_AF_UW_W','Q_placenta_W','Q_check_sum_W', ...
    'Q_skin_AF_UW_pct','Q_placenta_pct', ...
    'Q_UV_mLmin','Q_CVO_estimate_mLmin','Umbilical_pct_of_CVO'};

data_cb3 = [Tm_B_co', Tm_B_ax', Taf_sw, Tsk_sw, Ta_sw, ...
    dT_scalp_core_sw, dT_scalp_ax_sw, dT_af_core_sw, dT_af_ax_sw, ...
    repmat(Q_total,nB,1), Q_wall_sw, Q_plac_sw, Q_check_sw, ...
    Q_wall_pct, Q_plac_pct, ...
    repmat(Q_UV_mLmin_val,nB,1), repmat(Q_CVO_mLmin_val,nB,1), ...
    repmat(umb_pct_val,nB,1)];

T_cb3 = array2table(data_cb3, 'VariableNames', cb3_hdr);
writetable(T_cb3, fname_xls, 'Sheet', 'Checks_and_Balances');

fprintf('Excel saved → %s\n', fname_xls);
fprintf('  Sheet 1: single point at T_m = 37 °C  (%d rows)\n', height(T_sp));
fprintf('  Sheet 2: sweep T_m = %.1f–%.1f °C  (%d rows)\n', Tm_B_co(1), Tm_B_co(end), nB);

[~, idx37] = min(abs(Tm_B_co - 37.0));   % nearest to 37 °C (avoids float equality)
fprintf('\n--- Checks & Balances (T_m = 37 °C) ---\n');
fprintf('  Q_BMR total           = %.4f W\n', Q_total);
fprintf('  Q via skin/AF/UW      = %.4f W  (%.1f%%)\n', Q_wall_sw(idx37), Q_wall_pct(idx37));
fprintf('  Q via placenta        = %.4f W  (%.1f%%)\n', Q_plac_sw(idx37), Q_plac_pct(idx37));
fprintf('  Energy check (sum)    = %.4f W  (error = %.2e W)\n', ...
    Q_check_sw(idx37), Q_check_sw(idx37) - Q_total);
fprintf('  Q_UV                  = %.1f mL/min\n', Q_UV_mLmin_val);
fprintf('  Q_CVO estimate        = %.1f mL/min\n', Q_CVO_mLmin_val);
fprintf('  Umbilical / CVO       = %.1f%%\n', umb_pct_val);
fprintf('  dT scalp - T_core     = %.4f °C\n', Tsk_sw(idx37) - 37.0);
fprintf('  dT scalp - T_axillary = %.4f °C\n', Tsk_sw(idx37) - (37.0 - dT_ax));
fprintf('  dT T_af - T_core      = %.4f °C\n', Taf_sw(idx37) - 37.0);
fprintf('  dT T_af - T_axillary  = %.4f °C\n', Taf_sw(idx37) - (37.0 - dT_ax));


%% ======================================================================
%  LOCAL FUNCTIONS  (must appear after all script statements — MATLAB R2016b+)
%% ======================================================================

function out = solve_26x26(Tmother, ...
        Nseg, Nlay, Phi, TC_cm, TC_mf, TC_fs, ...
        h_conv, A_surf, qdot_co, qdot_mu, qdot_fa, qdot_sk, ...
        V_lay, h_uterus, A_uterus, umb)
    %SOLVE_26X26  Assemble and solve the 26×26 steady-state fetal system.
    %
    %  Unknowns x(1..24): x(4i-3)=T_co, x(4i-2)=T_mu, x(4i-1)=T_fa, x(4i)=T_sk
```

```
%                     x(25) = T_a  (central arterial blood pool)
%                     x(26) = T_af (amniotic fluid)
%
%  Node energy balance (each layer, each segment):
%    qdot·V  +  TC_in·(T_in − T)  −  TC_out·(T − T_out)
%            +  Φ·(T_a − T)  [+  h·A·(T_af − T_sk) for skin only]  = 0
%
%  T_a node:  Σ_layers Φ·T_layer − (Σ Φ + umb)·T_a = −umb·T_m
%  T_af node: Σ_segs h·A·T_sk  − (Σ h·A + h_u·A_u)·T_af = −h_u·A_u·T_m

N    = Nseg*Nlay + 2;   % 26
iTA  = N - 1;            % 25  T_a
iTAF = N;                % 26  T_af

A_mat = zeros(N, N);
b_vec = zeros(N, 1);

for i = 1:Nseg
    ci = 4*(i-1)+1;  mi = 4*(i-1)+2;
    fi = 4*(i-1)+3;  si = 4*(i-1)+4;

    % -- Core -----------------------------------------------------------
    A_mat(ci,ci)  = -(TC_cm(i) + Phi(i,1));
    A_mat(ci,mi)  =   TC_cm(i);
    A_mat(ci,iTA) =   Phi(i,1);
    b_vec(ci)     = - qdot_co(i)*V_lay(i,1);

    % -- Muscle ---------------------------------------------------------
    A_mat(mi,ci)  =   TC_cm(i);
    A_mat(mi,mi)  = -(TC_cm(i) + TC_mf(i) + Phi(i,2));
    A_mat(mi,fi)  =   TC_mf(i);
    A_mat(mi,iTA) =   Phi(i,2);
    b_vec(mi)     = - qdot_mu(i)*V_lay(i,2);

    % -- Fat ------------------------------------------------------------
    A_mat(fi,mi)  =   TC_mf(i);
    A_mat(fi,fi)  = -(TC_mf(i) + TC_fs(i) + Phi(i,3));
    A_mat(fi,si)  =   TC_fs(i);
    A_mat(fi,iTA) =   Phi(i,3);
    b_vec(fi)     = - qdot_fa(i)*V_lay(i,3);

    % -- Skin -----------------------------------------------------------
    A_mat(si,fi)   =   TC_fs(i);
    A_mat(si,si)   = -(TC_fs(i) + h_conv(i)*A_surf(i) + Phi(i,4));
    A_mat(si,iTAF) =   h_conv(i)*A_surf(i);
    A_mat(si,iTA)  =   Phi(i,4);
    b_vec(si)      = - qdot_sk(i)*V_lay(i,4);
end

% -- T_a row ------------------------------------------------------------
for i = 1:Nseg
    for l = 1:Nlay
        A_mat(iTA, 4*(i-1)+l) = A_mat(iTA, 4*(i-1)+l) - Phi(i,l);
    end
end
A_mat(iTA,iTA) = sum(Phi(:)) + umb;
b_vec(iTA)     = umb * Tmother;

% -- T_af row -----------------------------------------------------------
for i = 1:Nseg
    A_mat(iTAF, 4*i) = h_conv(i)*A_surf(i);
end
A_mat(iTAF,iTAF) = -(sum(h_conv.*A_surf) + h_uterus*A_uterus);
b_vec(iTAF)      = -h_uterus*A_uterus*Tmother;

% -- Solve --------------------------------------------------------------
Temps = A_mat \ b_vec;

out.Ta    = Temps(iTA);
```

```
    out.Taf   = Temps(iTAF);
    out.T_co  = Temps(1:4:4*Nseg-3);
    out.T_mu  = Temps(2:4:4*Nseg-2);
    out.T_fa  = Temps(3:4:4*Nseg-1);
    out.T_sk  = Temps(4:4:4*Nseg);
    out.Temps = Temps;
end


function apply_pubstyle(hFig, width_in, height_in)
    %APPLY_PUBSTYLE  Publication format + explicit white/black colours.
    %  Forces white backgrounds and black text/lines on every axes and text
    %  object, defeating MATLAB dark-theme overrides that survive groot resets.
    if nargin < 2, width_in  = 3.5; end
    if nargin < 3, height_in = 2.0; end

    % -- Figure ------------------------------------------------------------
    set(hFig, 'Color', [1 1 1]);   % white figure window background
    set(hFig, 'Units',             'inches', ...
              'Position',          [1 1 width_in height_in], ...
              'PaperUnits',        'inches', ...
              'PaperPosition',     [0 0 width_in height_in], ...
              'PaperSize',         [width_in height_in], ...
              'PaperPositionMode', 'manual', ...
              'Renderer',          'painters');

    % -- Axes --------------------------------------------------------------
    ax = findall(hFig, 'type', 'axes');
    for k = 1:numel(ax)
        ax(k).Color           = [1 1 1];           % axes plot area: white
        ax(k).XColor          = [0 0 0];           % x spine, ticks, labels
        ax(k).YColor          = [0 0 0];           % y spine, ticks, labels
        ax(k).ZColor          = [0 0 0];
        ax(k).GridColor       = [0.85 0.85 0.85];
        ax(k).MinorGridColor  = [0.92 0.92 0.92];
        ax(k).FontName        = 'Arial';
        ax(k).FontSize        = 9;
        ax(k).LineWidth       = 1;
        ax(k).Box             = 'on';
        ax(k).TickDir         = 'out';
        % Force tick label and title colours (dark theme sets these independently)
        ax(k).XLabel.Color    = [0 0 0];
        ax(k).YLabel.Color    = [0 0 0];
        ax(k).Title.Color     = [0 0 0];
        try ax(k).LooseInset = ax(k).TightInset; end
    end

    % -- All text objects (titles, annotations, colorbars) -----------------
    txt = findall(hFig, 'type', 'text');
    for k = 1:numel(txt)
        txt(k).Color    = [0 0 0];
        txt(k).FontName = 'Arial';
    end

    % -- Legends -----------------------------------------------------------
    lg = findall(hFig, 'type', 'legend');
    for k = 1:numel(lg)
        lg(k).FontName  = 'Arial';
        lg(k).FontSize  = 9;
        lg(k).Box       = 'off';
        lg(k).Color     = [1 1 1];   % white legend background
        lg(k).TextColor = [0 0 0];   % black legend text
    end
end
```

### S.11 Lavesson et al. maternal axillary vs. fetal scalp temperatures for model validation

**Table 6.** Lavesson et al. [57,58] data used in validation of the two simulations. Save as lavesson_full_data.xlsx in same folder as above codes in order to generate Figure 2a.

| Maternal Axillary Temp [degC] | Fetal Scalp Temp[degC] |
|---|---|
| 36.82 | 36.95 |
| 35.39 | 37.42 |
| 35.28 | 37.41 |
| 35.29 | 37.12 |
| 34.82 | 36.57 |
| 35.66 | 36.46 |
| 35.90 | 36.39 |
| 36.05 | 36.47 |
| 35.95 | 36.69 |
| 35.72 | 36.70 |
| 36.53 | 36.74 |
| 36.25 | 36.85 |
| 36.12 | 37.13 |
| 36.24 | 37.11 |
| 36.24 | 37.11 |
| 35.80 | 37.14 |
| 35.61 | 37.64 |
| 35.76 | 37.61 |
| 35.28 | 38.37 |
| 35.81 | 38.44 |
| 35.97 | 38.98 |
| 36.05 | 39.24 |
| 36.14 | 38.36 |
| 36.35 | 38.28 |
| 36.45 | 38.44 |
| 36.78 | 38.34 |
| 36.55 | 38.00 |
| 36.57 | 37.83 |
| 36.57 | 37.84 |
| 36.54 | 37.91 |
| 36.66 | 37.82 |
| 36.66 | 37.91 |
| 36.25 | 37.76 |
| 36.27 | 37.68 |
| 36.12 | 37.61 |
| 36.05 | 37.52 |
| 36.12 | 37.51 |

| | |
|---|---|
| 36.07 | 37.45 |
| 36.06 | 37.28 |
| 36.18 | 37.29 |
| 36.18 | 37.40 |
| 36.21 | 37.48 |
| 36.28 | 37.40 |
| 36.39 | 37.35 |
| 36.34 | 37.03 |
| 36.52 | 36.95 |
| 36.56 | 37.07 |
| 36.65 | 36.99 |
| 36.72 | 36.96 |
| 36.59 | 37.18 |
| 36.67 | 37.09 |
| 37.29 | 37.42 |
| 36.68 | 37.25 |
| 36.70 | 37.32 |
| 36.73 | 37.18 |
| 36.79 | 37.13 |
| 36.85 | 37.21 |
| 36.89 | 37.13 |
| 36.99 | 37.16 |
| 36.96 | 37.28 |
| 36.96 | 37.22 |
| 36.96 | 37.22 |
| 37.01 | 37.37 |
| 37.01 | 37.37 |
| 36.94 | 37.37 |
| 37.05 | 37.29 |
| 36.42 | 37.61 |
| 36.37 | 37.63 |
| 36.38 | 37.63 |
| 36.47 | 37.72 |
| 36.55 | 37.70 |
| 36.49 | 37.57 |
| 36.54 | 37.49 |
| 36.47 | 37.48 |
| 36.47 | 37.48 |
| 36.58 | 37.38 |
| 36.59 | 37.60 |
| 36.69 | 37.59 |

| | |
|---|---|
| 36.68 | 37.51 |
| 36.67 | 37.43 |
| 36.76 | 37.77 |
| 36.76 | 37.77 |
| 36.69 | 37.71 |
| 36.77 | 37.69 |
| 36.82 | 37.46 |
| 36.83 | 37.37 |
| 36.83 | 37.37 |
| 36.82 | 37.45 |
| 36.84 | 37.57 |
| 36.93 | 37.59 |
| 36.91 | 37.51 |
| 37.00 | 37.50 |
| 37.08 | 37.50 |
| 37.12 | 37.54 |
| 37.13 | 37.42 |
| 37.12 | 37.54 |
| 36.91 | 37.67 |
| 36.95 | 37.77 |
| 36.89 | 37.69 |
| 36.89 | 37.69 |
| 37.04 | 37.61 |
| 37.04 | 37.61 |
| 37.01 | 37.68 |
| 36.82 | 38.00 |
| 36.95 | 38.15 |
| 36.93 | 38.01 |
| 36.91 | 37.89 |
| 36.92 | 37.90 |
| 37.03 | 37.86 |
| 37.05 | 37.78 |
| 37.14 | 37.83 |
| 37.11 | 37.92 |
| 37.02 | 38.02 |
| 37.03 | 38.08 |
| 37.03 | 38.08 |
| 37.16 | 37.69 |
| 37.26 | 37.54 |
| 37.23 | 37.61 |
| 37.40 | 37.61 |

| | |
|---|---|
| 37.48 | 37.65 |
| 37.49 | 37.65 |
| 37.40 | 37.61 |
| 37.34 | 37.56 |
| 37.24 | 37.70 |
| 37.38 | 37.69 |
| 37.38 | 37.69 |
| 37.31 | 37.67 |
| 37.25 | 37.78 |
| 37.35 | 37.83 |
| 37.48 | 37.78 |
| 37.22 | 37.87 |
| 37.28 | 37.89 |
| 37.36 | 37.88 |
| 37.33 | 37.76 |
| 37.43 | 37.84 |
| 37.55 | 37.91 |
| 37.52 | 37.87 |
| 37.52 | 37.87 |
| 37.36 | 37.88 |
| 37.17 | 38.01 |
| 37.17 | 38.23 |
| 37.17 | 38.23 |
| 37.10 | 38.25 |
| 37.24 | 38.25 |
| 37.24 | 38.13 |
| 37.30 | 37.98 |
| 37.30 | 37.98 |
| 37.24 | 38.04 |
| 37.33 | 38.06 |
| 37.42 | 37.95 |
| 37.52 | 37.99 |
| 37.35 | 38.15 |
| 37.34 | 38.23 |
| 37.42 | 38.03 |
| 37.52 | 38.13 |
| 37.40 | 38.11 |
| 37.43 | 38.17 |
| 37.52 | 38.13 |
| 37.51 | 38.06 |
| 37.63 | 38.03 |

| | |
|---|---|
| 37.68 | 37.94 |
| 37.75 | 37.96 |
| 37.78 | 38.07 |
| 37.90 | 38.05 |
| 37.86 | 38.13 |
| 37.71 | 38.06 |
| 37.75 | 38.15 |
| 37.61 | 38.12 |
| 37.59 | 38.19 |
| 37.34 | 38.57 |
| 36.93 | 38.97 |
| 37.00 | 38.97 |
| 37.04 | 39.04 |
| 37.31 | 38.96 |
| 37.43 | 38.99 |
| 37.33 | 38.82 |
| 37.36 | 38.75 |
| 37.43 | 38.67 |
| 37.43 | 38.80 |
| 37.58 | 38.82 |
| 37.72 | 38.85 |
| 37.72 | 38.95 |
| 37.69 | 39.11 |
| 37.66 | 39.06 |
| 37.66 | 39.06 |
| 37.72 | 39.51 |
| 37.77 | 39.58 |
| 38.03 | 39.48 |
| 38.08 | 39.29 |
| 37.54 | 38.35 |
| 37.62 | 38.27 |
| 37.92 | 38.22 |
| 37.72 | 38.29 |
| 37.81 | 38.19 |
| 37.87 | 38.29 |
| 37.79 | 38.30 |
| 37.73 | 38.29 |
| 37.63 | 38.35 |
| 37.65 | 38.43 |
| 37.60 | 38.51 |
| 37.58 | 38.64 |

| | |
|---|---|
| 37.65 | 38.70 |
| 37.72 | 38.57 |
| 37.67 | 38.54 |
| 37.67 | 38.54 |
| 37.76 | 38.47 |
| 37.78 | 38.65 |
| 37.92 | 38.66 |
| 37.89 | 38.48 |
| 37.89 | 38.49 |
| 37.88 | 38.56 |
| 37.93 | 38.33 |
| 38.03 | 38.28 |
| 38.00 | 38.37 |
| 37.99 | 38.45 |
| 38.07 | 38.41 |
| 38.14 | 38.39 |
| 38.22 | 38.41 |
| 38.31 | 38.43 |
| 38.41 | 38.59 |
| 38.47 | 38.80 |
| 37.99 | 38.53 |
| 38.04 | 38.62 |
| 38.09 | 38.55 |
| 38.10 | 38.48 |
| 38.17 | 38.52 |
| 38.18 | 38.59 |
| 38.00 | 38.84 |
| 38.00 | 38.84 |
| 37.93 | 38.77 |
| 38.16 | 38.71 |
| 38.22 | 38.77 |
| 38.26 | 38.63 |
| 38.33 | 38.71 |
| 38.41 | 38.73 |
| 38.41 | 38.73 |
| 38.33 | 38.71 |
| 38.78 | 38.91 |
| 38.73 | 39.03 |
| 38.58 | 38.96 |
| 38.58 | 38.96 |
| 38.66 | 39.06 |

| | |
|---|---|
| 38.61 | 39.03 |
| 38.61 | 39.03 |
| 38.41 | 38.95 |
| 38.32 | 38.96 |
| 38.41 | 38.95 |
| 38.34 | 38.87 |
| 38.11 | 38.93 |
| 38.13 | 39.00 |
| 38.17 | 39.06 |
| 38.22 | 39.02 |
| 38.25 | 38.96 |
| 38.94 | 39.23 |
| 39.14 | 39.58 |
| 39.15 | 39.79 |
| 38.92 | 39.80 |
| 38.83 | 39.69 |
| 38.71 | 39.67 |
| 38.47 | 39.93 |
| 38.37 | 39.52 |
| 38.50 | 39.57 |
| 38.68 | 39.44 |
| 38.80 | 39.40 |
| 38.89 | 39.36 |
| 38.86 | 39.31 |
| 38.86 | 39.31 |
| 38.78 | 39.27 |
| 38.47 | 39.12 |
| 38.38 | 39.15 |
| 38.44 | 39.23 |
| 38.35 | 39.23 |
| 38.40 | 39.36 |
| 38.61 | 39.18 |
| 38.63 | 39.27 |
| 38.67 | 39.15 |
| 38.67 | 39.15 |
| 38.71 | 39.22 |

*Surgical and radiologic anatomy (English ed.)* **41**, 431–439. (doi:10.1007/s00276-018-2121-2)

52. Fiala D, Lomas KJ, Stohrer M. 1999 A computer model of human thermoregulation for a wide range of environmental conditions: the passive system. *J. Appl. Physiol.*
53. Sokoloff L. 1959 Circulation and metabolism of brain in relation to the process of aging. *The process of aging in the nervous system* , 113–126.
54. Gruenwald P. 1966 Growth of the human fetus: I. Normal growth and its variation. *Am. J. Obstet. Gynecol.* **94**, 1112–1119.
55. Rudolph AM. 1985 Distribution and regulation of blood flow in the fetal and neonatal lamb. *Circ. Res.* **57**, 811–821.
56. Gruenwald P, Minh HN. 1960 Evaluation of Body and Organ Weights in Perinatal Pathology: I. Normal Standards Derived from Autopsies. *Am. J. Clin. Pathol.* **34**, 247–253. (doi:10.1093/ajcp/34.3.247)
57. Lavesson T, Källén K, Olofsson P. 2018 Fetal and maternal temperatures during labor and delivery: a prospective descriptive study. *The Journal of Maternal-Fetal & Neonatal Medicine* **31**, 1533–1541.
58. Lavesson T. 2017 Continuous Intrapartum Maternal and Fetal Temperature Monitoring . Doctoral, Lunch University, Malmo, Sweden.
